# Hydralazine inhibits cysteamine dioxygenase to treat preeclampsia and senesce glioblastoma

**DOI:** 10.1101/2024.12.19.629450

**Authors:** Kyosuke Shishikura, Jiasong Li, Yiming Chen, Nate R. McKnight, Katelyn A. Bustin, Eric W. Barr, Snehil R. Chilkamari, Mahaa Ayub, Sun Woo Kim, Zongtao Lin, Ren-Ming Hu, Kelly Hicks, Xie Wang, Donald M. O’Rourke, J. Martin Bollinger, Zev A. Binder, William H. Parsons, Kirill A. Martemyanov, Aimin Liu, Megan L. Matthews

## Abstract

The vasodilator hydralazine (HYZ) has been used clinically for ∼ 70 years and remains on the World Health Organization’s List of Essential Medicines as a therapy for preeclampsia. Despite its longstanding use and the concomitant progress toward a general understanding of vasodilation, the target and mechanism of HYZ have remained unknown. We show that HYZ selectively targets 2-aminoethanethiol dioxygenase (ADO) by chelating its metal cofactor and alkylating one of its ligands. This covalent inactivation slows entry of proteins into the Cys/N-degron pathway that ADO initiates. HYZ’s capacity to stabilize regulators of G-protein signaling (RGS4/5) normally marked for degradation by ADO explains its effect on blood vessel tension and comports with prior associations of insufficient RGS levels with human preeclampsia and analogous symptoms in mice. The established importance of ADO in glioblastoma led us to test HYZ in these cell types. Indeed, a single treatment induced senescence, suggesting a potential new HYZ-based therapy for this deadly brain cancer.

## Main text

Hydralazine (HYZ, Apresoline®) is one of the oldest FDA-approved vasodilators. Originally developed as a treatment for malaria, it was introduced into the clinic more than 70 years ago. Although many other vasodilatory and antihypertensive drugs have been developed since that time, HYZ remains on the World Health Organization’s Essential Medicine List(*1*) as a treatment for (pre)eclampsia(*2*) and hypertensive crisis(*2, 3*). Despite decades of advancements in our general understanding of vasodilation physiology, HYZ’s mechanism of action is still unknown.

Vasodilation is an adaptive physiological response for immediate and preferential delivery of oxygen and essential nutrients to microenvironments consuming them at rates exceeding their delivery. When blood flow is impaired by increased vascular resistance of maladaptive vessels, blood pressure increases and hypertension can ensue. Dynamic constriction and dilation of vessels are modulated by changes in calcium (Ca^2+^) ion concentrations within vascular smooth muscle cells. As such, both normal and pharmacologically-induced vasodilation results from decreasing intracellular Ca^2+^ levels. The specific mechanisms to achieve this effect are well-established in general and vary for the different classes of vasodilators(*4*). For the case of HYZ, however, they remain unknown.

To date, a number of potential targets for HYZ have been proposed. They include prolyl hydroxylase domain (PHD) enzymes(*5*), the Keap1-Nrf2 complex(*6*), aldehyde oxidase (AO)(*7*), protein kinase A(*8*), and glutamine oxaloacetate transaminase 1 (GOT1)(*9*). Supporting evidence has consisted of either downstream physiological effects expected for inhibition of the proposed target or inhibition of that enzyme *in vitro*. In no case have both lines of evidence been provided to directly connect the engagement of the target by the drug to the cellular effects of its inhibition. Moreover, no links between inhibition and the effects on Ca^2+^ levels expected to be required to drive vasodilation have been established.

Previous work has shown that chemical probes bearing an organohydrazine reactive group can undergo covalent coupling to the active sites of enzymes that use electron-deficient cofactors in catalysis(*10*). In a subset of these coupling reactions, the hydrazine pharmacophore is lost (most likely as N_2_) and the appended carbon radical fragment couples to an amino acid side chain in the enzyme active site proximal to the initiating cofactor. HYZ, as a hydrazine-containing drug, could potentially react with electron-deficient catalytic cofactors by the same manifold (fig. S1).

## RESULTS

### HYZ selectively targets ADO in cells and tissues

To uncover the immediate target(s) of HYZ, we appended an alkyne functionality (propargyl ether group) at a position (C4) distal and undisruptive to the drug’s nitrogen-containing pharmacophore(s)(*11*) to create the chemical probe ‘HYZyne’ (Fig. 1A). The alkyne allows for covalent capture and identification of the posited enzyme adduct with the phthalazine fragment of HYZ left behind upon hydrazine activation and loss of N_2_. As expected, HYZyne-treated HEK293T cells showed concentration-dependent protein labelling (fig. S2A), following conjugation to a fluorophore (*e.g.,* rhodamine-azide) by Cu(I)-catalyzed azide-alkyne cycloaddition (CuAAC, or “click” chemistry) and visualization by SDS-PAGE in-gel fluorescent scanning(*11*). Tagging of cellular proteins was blocked by pretreatment (15 min, 37 °C) with HYZ (fig. S2B). Importantly, band intensity was not correlated with protein expression level (fig. S2, A and B). Next, we applied established chemical proteomics mass spectrometry-based approaches(*12*) to identify and quantify high-reactivity and high-stoichiometry targets of HYZ(yne)(*10, 13*). By contrast to our other hydrazine-containing probes and drugs that identified > 100 targets (externally validated) when analyzed under the same conditions(*10, 14*), HYZyne was surprisingly selective for a single enzyme target – 2-aminoethanethiol (i.e. cysteamine) dioxygenase (ADO) (Fig. 1B and Data S1). ADO, a Fe(II)-dependent thiol dioxygenase, oxidizes cysteine-derived thiol groups in both protein substrates and cysteamine using molecule oxygen, as elaborated below (Fig. 2A)(*15, 16*).

**Fig. 1.**
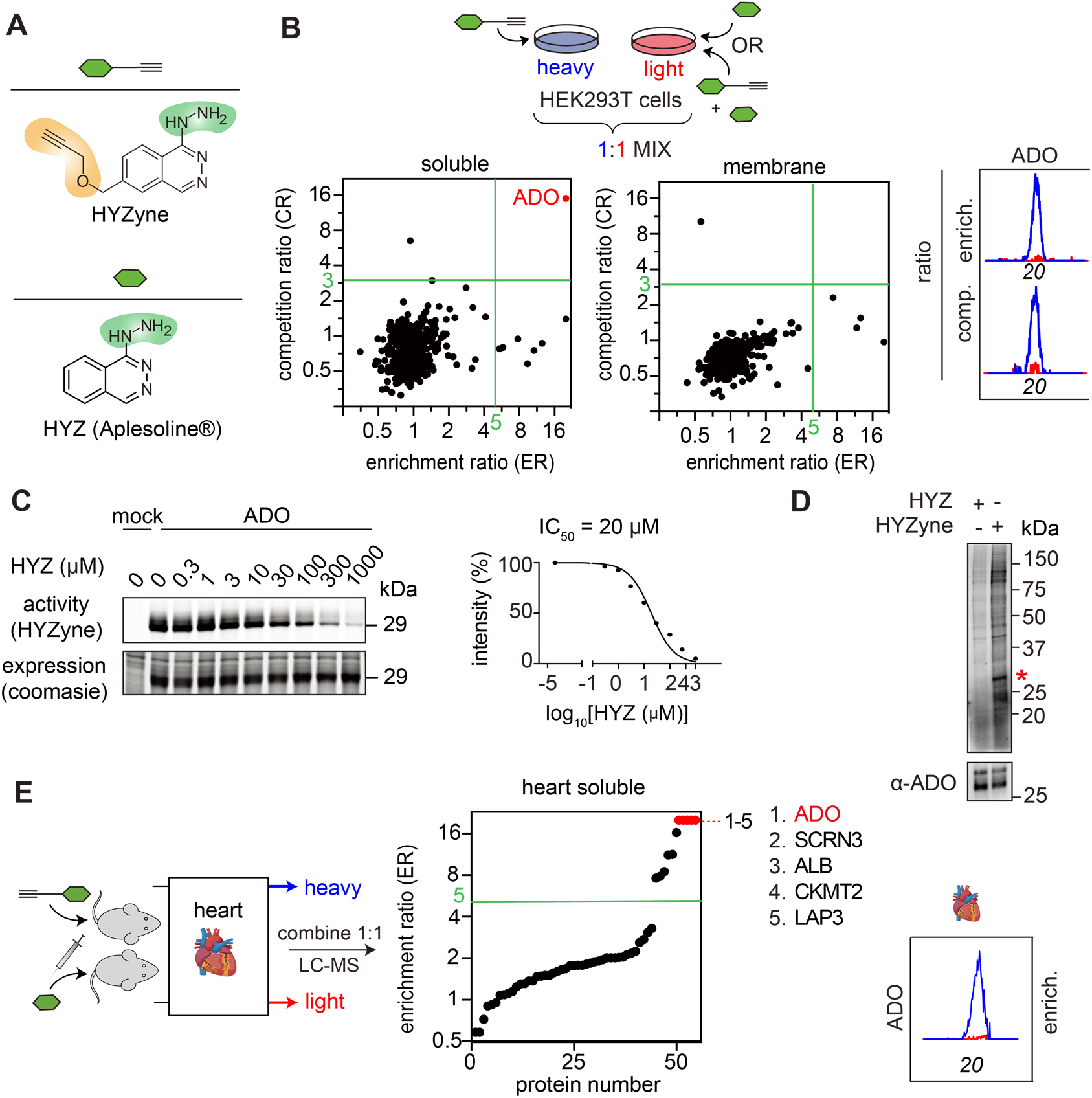
Identification of protein targets of HYZ. (**A**) Structures of hydralazine drug (HYZ) and the corresponding alkynylated drug probe (HYZyne). The reactive hydrazine pharmacophores are shown in green and chemical modifications made to the parent drug to incorporate the clickable alkyne handle are shown in orange. (**B**) Quadrant plot of average competition versus enrichment SILAC ratios from quantitative proteomics experiments (left). HYZyne-targeted proteins (upper right quadrant) are highlighted in red. Extracted MS1 chromatograms and corresponding heavy/light ratios (enrichment and competition) for representative tryptic peptides of endogenous high-reactivity targets. (**C**) HYZyne-labeling (upper) and coomasie staining (lower) for HYZyne-treated cells expressing ADO. The first lane in each panel corresponds to a control transfection (‘mock’) with the appropriate empty expression vector as described in Supplementary Methods. Treatment of HEK293T cells with HYZ blocks in a concentration-dependent manner the labelling of ADO by HYZyne (1 mM, 0.5 h). (**D**) Probe labeling (upper) and Western blot for ADO (lower) for soluble proteome from non-transfected HEK293T cells harvested 0.5 h post-treatment with 10 μM HYZyne. Asterisk (*) highlights a single band corresponding to ADO. (**E**) Quadrant plot of average enrichment ReDiMe ratios versus protein number for HYZyne from quantitative proteomics experiments in the soluble proteomes of mouse heart 4 h post-injection with HYZ or HYZyne (50 mg/kg, intraperitoneally). Protein targets with ER ≥ 5 were considered high-reactivity targets and are annotated.

**Fig. 2.**
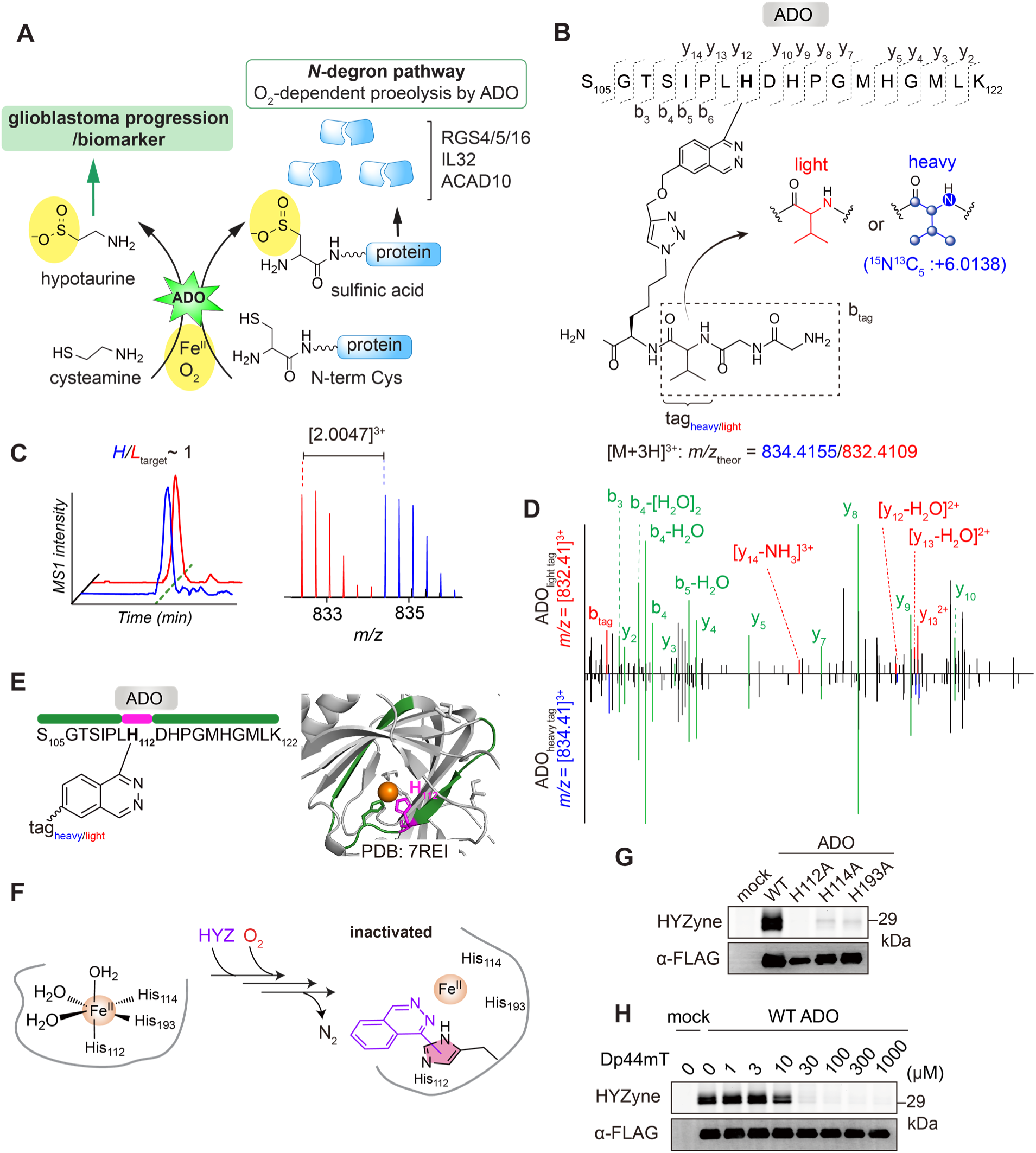
Covalent modification of active site histidine residues in ADO. (**A**) Scheme of catalytic machinery and function of ADO. ADO is an Fe(II)- and O_2_-dependent dioxygenase that oxidizes thiol group of cysteamine and *N-t*erminal Cys in protein substrates including regulator of G protein signaling (RGS4/5/16), interleukin-32 (IL32) and acyl-coA dehydrogenase family member 10 (ACAD10). Oxidized protein substrates are subjected to degradation. (**B**) Structures and theoretical parent masses of heavy- and light-tagged ADO peptides labeled by HYZyne and processed by the isoTOP-ABPP method(*19*). (**C**) Extracted triply charged MS1 ion chromatograms (left) and corresponding isotopic envelopes (right) for co-eluting heavy- and light-tagged peptides labelled by HYZyne (blue and red, respectively). (**D**) Comparison of high-resolution MS2 spectra generated from light-versus heavy-tagged parent ions for the ADO peptide pair in C. Unshifted ions are shown in green and the shifted ion series in blue (heavy) and red (light). (**E**) Labeled peptide in the ADO crystal structure (PDB: 7REI). The identity of the HYZyne labeled peptides (aa105–122) and site (H112) are shown in green and magenta, respectively. Metal is colored orange. (**F**) Reaction scheme and MS results-based prediction of the structure of the HYZyne. (**G**) 100 μM HYZyne labeling of wild-type ADO but not catalytically inactive variants. Probe labeling (upper) and expression profiles (lower) for probe-treated cells overexpressing the indicated protein target or inactive variant thereof. (**H**) Dose-dependent inhibition of HYZyne-labeling with iron chelator (Dp44mT). Expression profiles of overexpressed wild-type enzymes were visualized by Western blot.

Because HYZ is also effective in preclinical mouse models of hypertension, we repeated the same target identification experiments in two mouse cell lines [RAW 264.7 macrophage and Neuro 2a neuroblastoma cells (fig. S3 and Data S2)]. Consistent with the HYZyne target profile from HEK293T cells, mouse ADO was identified as a primary target of HYZ in both cell types (fig. S3 and Data S2). We next validated that HYZ reactivity towards endogenous ADO was conserved in HYZ(yne)-treated cells transiently overexpressing *C-*terminally FLAG-tagged ADO. HYZyne-labelled protein bands were competed by HYZ in a concentration-dependent manner; quantitative analysis of the intensity of the ADO-associated 29 kDa band furnished an initial *in situ* IC_50_ ∼ 20 μM (Fig. 1C). To ensure that endogenous ADO activity could also be robustly detected at the calculated IC_50_ for recombinant ADO, *in situ* gel-based analysis was performed on HYZyne-treated (10 μM) HEK293T cells (Fig. 1D). Consistent with the proteomic profiling data, superior selectivity for endogenous ADO could be visualized in HEK293T cells [and at 100× lower concentrations than initial profiling experiments (Fig. 1D)], as illustrated by the single fluorescent band at 29 kDa.

To directly evaluate HYZ-targeted proteins in heart tissue and given the expected antihypertensive activity to be among the most clinically relevant tissues, wild-type mice were systemically treated with HYZ/HYZyne. Protein target enrichment was determined by comparing heart tissue proteomes from probe-treated (50 mg/kg, intraperitoneally, 4 h) *versus* drug-treated (50 mg/kg, intraperitoneally, 4 h) control mice. HYZyne-labelled targets were then identified by CuAAC coupling to a biotin-azide tag, streptavidin enrichment, and quantitative LC-MS using reductive dimethylation (ReDiMe)(*13*) chemistry to append isotopically “heavy” and “light” methyl groups to tryptic peptides of labelled proteins from probe-treated *versus* control-treated mice. Heavy/light protein enrichment ratios were determined by the median ratio of ≥2 unique quantified peptides per protein from soluble and membrane proteomes from heart tissue (Fig. 1E, fig. S4 and Data S2). The data confirmed that ADO is indeed an enriched protein target of HYZ in heart tissue following systemic delivery to the animal. In summary, ADO is the prevailing if not only target of HYZ across native biological systems tested including murine heart, and our global proteome-wide profiling data furnished ADO as the most likely pharmacological target of HYZ, retaining nearly complete selectivity for ADO across three orders of magnitude (0.01–1 mM). Consistent with this pharmacological mechanism of action, single nucleotide polymorphisms (SNP) that cause missense mutations in ADO (rs10995311: C to G; Pro39Ala) are associated with decreased blood pressure(*17, 18*).

### Cofactor coordination confers selective inactivation by HYZ

To define the site and reaction manifold for covalent capture of ADO by HYZ, we isolated ADO-specific peptide(s) modified by HYZyne from cells. This chemoproteomic approach is well-established(*19*), and leverages isotopically differentiated, protease-cleavable biotin-azide tags to enrich and release probe-labelled peptides that migrate and co-elute as mass differentiated pairs (Fig. 2, B and C). Among 7 ADO-specific peptides, that were identified with comparable MS1 intensities, only one contained a residue (His112) known to be essential for enzyme activity (Table S1)(*20*). This peptide pair was manually and computationally assigned as a triply-charged peptide comprising residues 105 to 122 of ADO according to the high-resolution MS2 spectra of y- and b-ions (Fig. 2D). Importantly, identification of the y_12_- and y_10_-ions resolved the site of probe capture between Leu111 and His112 (Fig. 2D), where His112 is one of ADO’s three cofactor ligand side chains that bind the enzyme’s metal center, a ferrous ion [Fe(II)], and are essential for catalysis (Fig. 2E, fig. S5 and Table S1)(*20*). The parent masses for the His112-containing peptide pair confer the radical manifold, as they were detected within 1 ppm error of the predicted product mass for arylation of the His112-coordinated side chain with the loss of hydrazine (–NHNH_2_) in the coupling event (Fig. 2F, fig. S5 and Table S1).

To confirm HYZ is selective for the active, functional resting state of the enzyme, we showed that HYZyne-labeling of ADO was abolished by substitution of active site residues (His112, His114 and His193) essential for metallocofactor binding and enzyme activity (Fig. 2G). By contrast, corresponding Cys-to-Ala mutants (Cys18, Cys132, Cys183, Cys220, Cys249 and Cys258) identified as other HYZyne-modified sites by chemoproteomics (Table S1) did not significantly impact the band intensity of HYZyne-labeled ADO (fig. S6). This data is consistent with substoichiometric labeling events that are nonessential for HYZ inhibitory activity, but that generated tryptic peptides with superior ionization efficiency during MS-based detection. Finally, pretreating cells with Dp44mT (0–1000 μM, 1 h), an iron chelator with reported specificity for ADO-bound Fe(II)(*21*) and that has structural homology to HYZ, abolished HYZyne labeling in a concentration-dependent manner (Fig. 2H). Collectively, the data show that HYZ is an enzyme-activated irreversible inhibitor of ADO. As such, HYZyne is also a fully functional and validated activity-based probe for ADO that can be used in the future to investigate how ADO activity is (dys)regulated and regulatory across biological contexts, for example, upon acute changes within a (pre/sub)hypoxic cellular environment(*21, 22*).

To capture the initial state of the HYZ•ADO reaction *in vitro* prior to reaction with O_2_, we first purified and crystalized the catalytically-competent cobalt-substituted human enzyme [Co(II)•ADO] as previously reported(*23, 24*). Soaking the resultant [Co(II)•ADO] crystals with excess HYZ resulted in the formation of the HYZ•Co(II)•ADO complex at a resolution of 1.88 Å. Electron density for the parent drug of HYZ was resolved to the active site metal center (Fig. 3A), while a second molecule was bound to the protein surface (fig. S7, and Table S2). The data refinement is summarized in fig. S8.

**Fig. 3.**
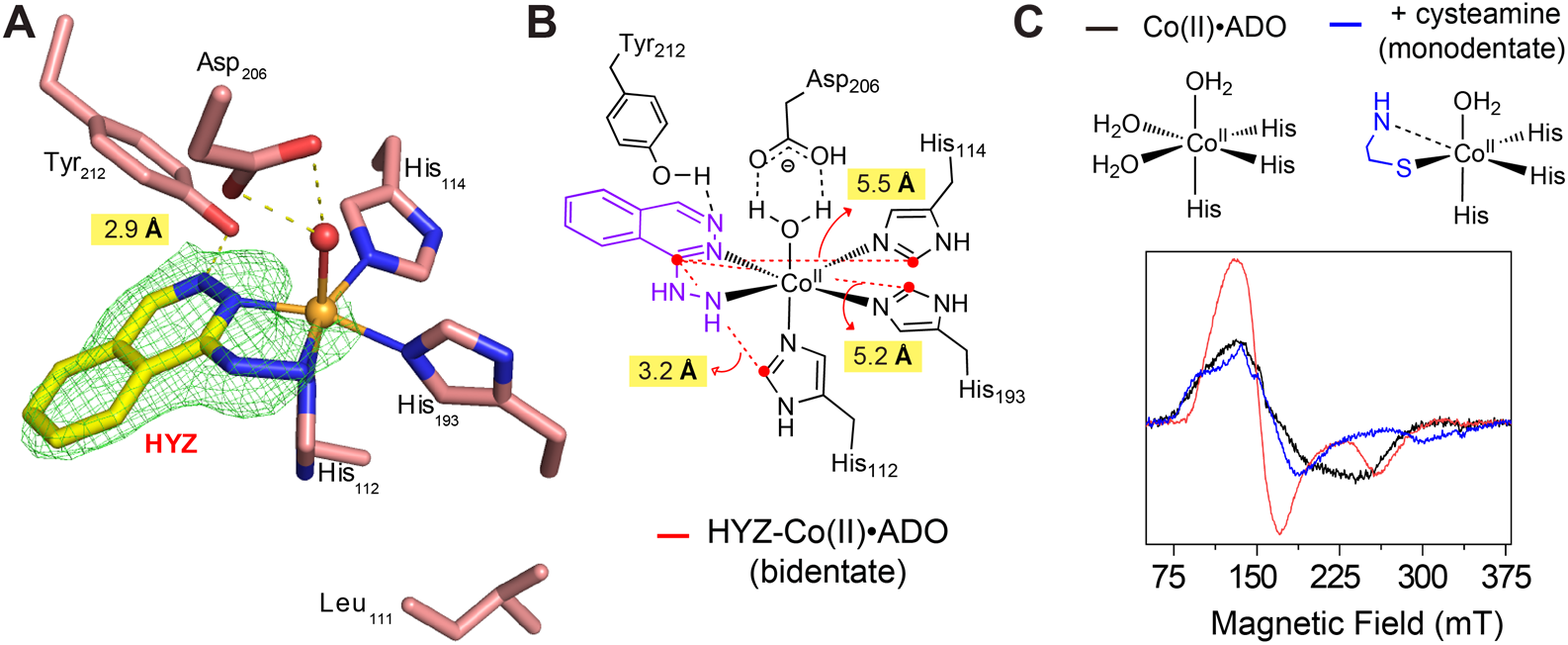
Determination of co-crystal structures of human ADO and HYZ. (**A**) Active site architecture of Co(II)•ADO. The X-ray crystal structure of HYZ-bound Co(II)•ADO complex was determined at 1.88 Å resolution. The *F_o_–F_c_* maps are contoured at 3.0 σ and colored in green. (**B**) A two-dimensional diagram of the active site showing the distances between HYZ and three histidine residues that coordinate to Co(II) center. (**C**) Schematic representation of the active site architecture of Co(II)•ADO (upper) and EPR spectra (lower) of 300 μM Co(II)•ADO (black) and upon adding 5 mM cysteamine (blue) or 1 mM HYZ (red) anaerobically. EPR measurements involved four scans at 3.17 mW microwave power at 30 K. Dashed line indicates potential interaction, not a covalent bond.

As expected, HYZ binds in a bidentate chelating mode to the same site as the primary substrate, replacing the two H_2_O molecules coordinated by His114 and His193 ligands. Additionally, a hydrogen bond between Tyr212 and the N3 atom of the phthalazine ring reveals a pseudo-tridentate interaction, underscoring the critical role of the phthalazine ring’s amine groups (N2 and N3) and the hydrazine moiety. These features likely represent non-modifiable elements in designing future ADO inhibitors that exploit this mechanism. Notably, the third H_2_O ligand in the equatorial position remains intact, stabilized by a hydrogen bond with the carboxylate group of Asp206. This structural evidence supports the classification of HYZ as a competitive inhibitor that preserves the enzyme’s capability to bind and activate O_2_, analogous to its interaction with endogenous substrate.

Further structural analysis revealed that His112 is significantly closer to HYZ (3.2 Å) compared to other ligands including His114 and His193 (5.5 and 5.2 Å, respectively) (Fig. 3B), explaining its preferential coupling in cellular assays. Conversely, Leu111, which projects away from the active site, is unlikely to participate in labeling (Fig. 3A). Alignment of Co(II)•ADO with and without HYZ confirmed that HYZ displaces the two equatorial H_2_O molecules without perturbing the active site architecture (fig. S9).

To further investigate the binding dynamics, electron paramagnetic resonance (EPR) spectroscopy was employed to compare the electronic environments of Co(II)•ADO bound to cysteamine versus HYZ. Cysteamine, which binds monodentately via its thiol group(*25*), caused a shift in *g*-tensor values. By contrast, the EPR spectrum of HYZ•Co(II)•ADO showed greater perturbation, consistent with HYZ’s bidentate chelation mode versus cysteamine (Fig. 3C and Table S3).

Finally, to confirm generation of the arylated product observed in cells, we reconstituted the reaction *in vitro* by monitoring O_2_ consumption under steady-state conditions with Fe(II)•ADO in the absence and presence of HYZ and O₂. Kinetic analysis, illustrated in a Lineweaver-Burk plot (fig. S10), revealed a near-pure competitive inhibition mode. Although the reciprocal plots do not converge perfectly at the y-axis, the deviation is minimal, and the *K*_i_ values calculated by global fitting the double reciprocal plots were 2.48 ± 0.42 and 13.88 ± 7.51 µM, respectively. This finding strongly supports HYZ’s predominant role as a competitive inhibitor. The result is consistent with the structural observation of bidentate binding of HYZ to the metal center (Fig. 3A) and the isolation of the arylated product in cells (Fig. 2).

### Accumulation of RGS4 and RGS5 upon ADO inhibition by HYZ

ADO is an evolutionarily-conserved iron-dependent thiol dioxygenase and enzymatic oxygen sensor that serves as the cell’s front-line responder to acute hypoxia(*22*). As the lowest affinity (highest-K_M_ for O_2_) oxygen-dependent enzyme known to date, ADO converts the metabolite cysteamine to hypotaurine and transduces the oxygen-dependent degradation of important regulatory proteins with amino-terminal cysteine residues. HYZ prevents formation of the *N*- degron recognition element – oxidation of *N*-Cys-SH groups to their corresponding sulfinic acids (-SO_2_) (Fig. 2A) – the essential post-translational modification required for targeted protein degradation by the ubiquitin-proteosome system(*26*). Known *N*-degron substrates of ADO include: i) three regulators of G protein signaling (RGS) 4/5/16, which are GTPase-activating proteins for heterotrimeric G proteins limiting duration and extent of GPCR signaling(*26–28*); ii) the angiogenic cytokine interleukin-32 (IL32)(*16*); and iii) the brain-specific metabolic enzyme acyl-CoA dehydrogenase family member 10 (ACAD10) central to beta-oxidation of fatty acids in the mitochondria(*29*). ADO’s oxidative post-translation modification serves as the essential substrate recognition element for arginyltransferases (*e.g.,* ATE1) that then append this destabilizing residue to the oxidized *N-*terminus, thereby promoting its targeted degradation (fig. S11).

To further test the impact of HYZ on ADO’s activity at the cellular level, we treated SH-SY5Y cells, a human neuroblastoma cell line derived from a metastatic bone tumor, commonly used for neuroscience research and previously used to study the impact of genetic deletion of ADO(*16, 21*). Among ADO substrates, we initially focused on RGS4 and RGS5 due to their established role in GPCR-mediated Ca^2+^ signaling and their clear connection to vasodilation(*30, 31*). Accordingly, in the absence of HYZ, RGS5 protein was undetectable, consistent with its transient expression and immediate ADO-catalyzed oxidation and subsequent degradation. By contrast, in the presence of HYZ (10 µM), RGS5 protein levels accumulated in < 1 h, which is consistent with the immediate blood-pressure-lowering effect observed in patients following administration of the drug (Fig. 4A)(*4*). HYZ inhibited targeted degradation of RGS5, with a half-maximal recovery of the protein with 1 μM, and maximum levels were observed with 10 μM (Fig. 4B and fig. S12). These results indicate that the pharmacological action of HYZ involves stable accumulation of RGS5. Similarly, RGS4 protein levels were upregulated with a half-maximal concentration of ∼ 1 μM (Fig. 4B and fig. S12). Notably, the effects of HYZ were specific to RGS proteins, as we did not observe any changes in ADO (Fig. 4B). To determine if the phenomenon was conserved across cell lines, we applied HYZ to HEK293T cells. While endogenous RGS4/5 levels were undetectable in HEK293T cells following treatment, endogenous RGS16, another known ADO protein substrate, was upregulated in response to HYZ (fig. S13A)(*32*). However, transient transfection of RGS5 into HEK293T cells allowed its detection and HYZ treatment led to a substantial increase in its levels by inhibiting both endogenous and overexpressed ADO in HEK293T cells (fig. S13, B and C). The data suggests the pharmacological effects of HYZ are universal across cell types.

**Fig. 4.**
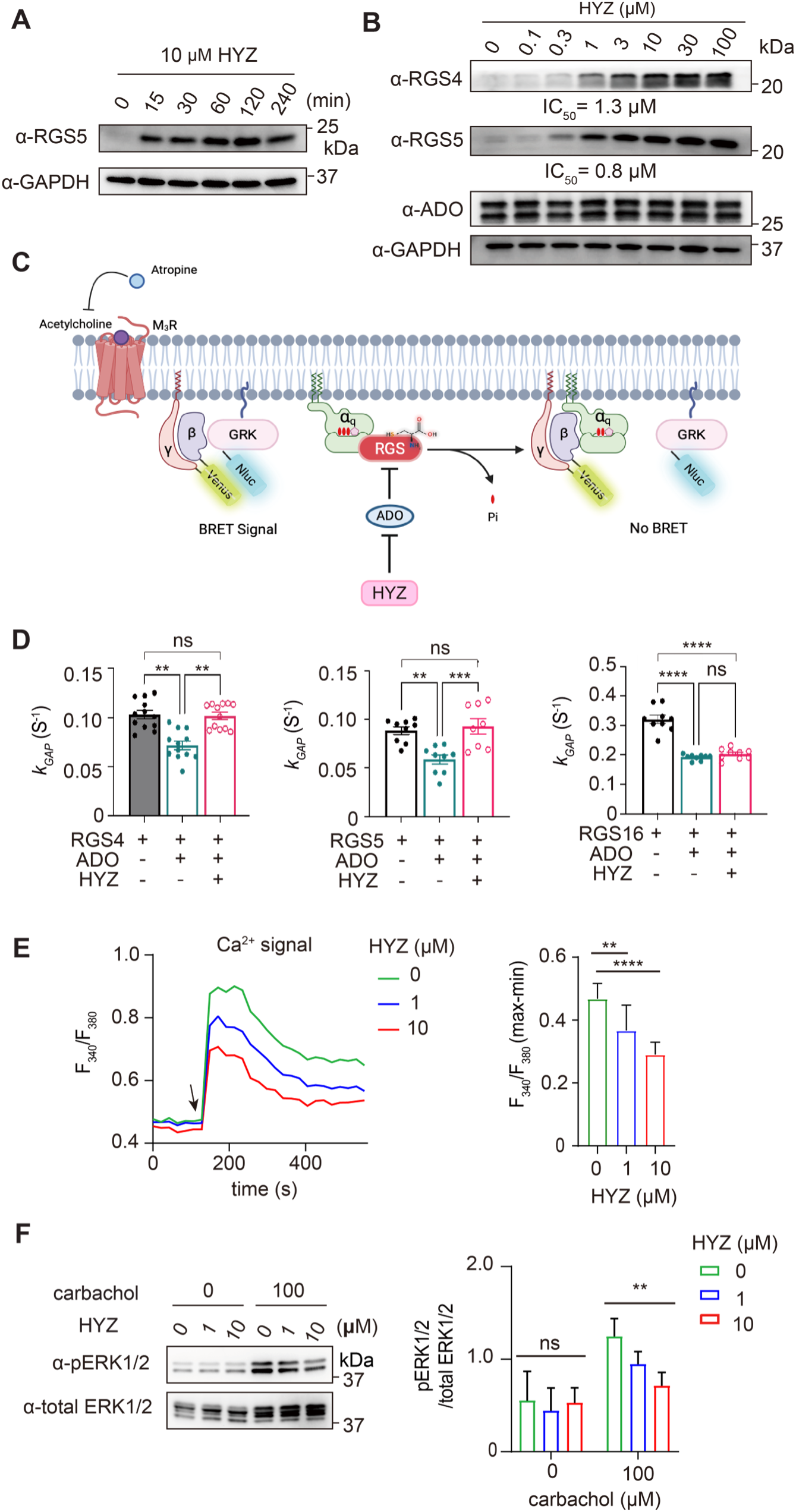
HYZ controls the ADO-dependent *N-*degron pathway and GPCR signaling in cells. (**A**) Time-dependent endogenous RGS5 and GAPDH protein levels in SH-SY5Y cells with 10 μM HYZ determined by Western blot. (**B**) Concentration-dependent endogenous RGS4/5, ADO, and GAPDH protein levels in SH-SY5Y cells after 1 h treatment of HYZ (0–100 μM) determined by Western blot. (**C**) Schematic diagram of the BRET assay. The rate of G protein deactivation is reflected as the rate of decrease in BRET signal. [Part of figure created with BioRender.com] (**D**) BRET assay assessing the effect of HYZ on *k*_GAP_ of RGS4/5/16. The relative activity *k*_GAP_ of a given RGS is calculated by subtraction of baseline deactivation rate from deactivation rate in the presence of RGS. For RGS4 (left) and RGS5 (middle), HYZ treatment (200 μM) restores the *k*_GAP_ in cells co-transfected with RGS and ADO back to the same level as cells transfected with RGS only. *k*_GAP_ is calculated from at least n = 3 independent experiments with each condition in triplicate. Data represent mean ± SEM. One-way ANOVA with Tukey’s multiple comparisons test, *\*\*p* < 0.005, *\*\*\*p* = 0.0008, *\*\*\*\*p* < 0.0001. ns; not significant. (**E**) Ca^2+^ influx in Fura-2 loaded SH- SY5Y cells after 1 h HYZ treatment (0–100 μM). Average traces of F_340_/F_380_ (let) and quantification of the maximum increase (right) in Ca^2+^ in response to 100 μM carbachol (127 s; arrow) normalized to minimum baseline Ca^2+^ levels. n = 7–8, mean ± SD. \*\**p* < 0.01, \*\*\*\**p* < 0.0001, one-way ANOVA with Tukey’s HSD. (**F**) Representative Western blot (left) and quantification (right) for total ERK1/2 and phosphorylated ERK1/2 (pERK1/2) in SH-SY5Y cells following exposure to HYZ (0–100 μM) for 1 h. Cells were incubated in the presence or absence of 100 μM carbachol for 5 min before harvest. n = 4, mean ± SD. Each column was tested against HYZ untreated control with one-way-ANOVA, followed by Tukey’s HSD. *\*\*p* < 0.01, ns; not significant.

### HYZ-promoted RGS accumulation attenuates GPCR signaling

Given that RGS proteins attenuate GPCR signaling, the demonstration of their accumulation upon treatment with HYZ provides a plausible link for the control of Ca^2+^-mediated signaling that drives vasodilation. As such, we evaluated the ADO- and HYZ-dependent changes on the rate of deactivation of G_αq_, the G protein pivotal to initiating Ca^2+^ release that RGS4/5/16 are known to regulate(*32*). Using an established bioluminescence resonance energy transfer (BRET)-based assay, we quantified the GTPase-activating protein (GAP) activity of RGS4, RGS5 and RGS16 by monitoring its ability to accelerate the rate of G_αq_-reassociation with its G_βγ_ subunits upon termination of GPCR signaling (*k*_GAP_; Fig. 4C)(*32*). First, we found that overexpression of ADO reduced RGS4 and RGS5 GAP activity upon inactivation of the muscarinic receptor (M_3_R), and that this effect was reversed by HYZ (200 µM) (Fig. 4D and fig. S14). The data indicate that HYZ indeed inhibits GPCR signaling via the ADO-RGS axis in a concentration-dependent manner (fig. S15). Notably, RGS16 GAP activity was not recovered under these conditions (Fig. 4D), suggesting that this RGS protein may be subject to other regulatory mechanisms that influence its ability to control M_3_R signaling. The complete dataset of GAP activity responses by HYZ is shown in fig. S16.

### HYZ reduces GPCR-mediated intracellular Ca^2+^ concentration

As G_αq_-coupled receptors mediate intracellular Ca^2+^ release in vascular smooth muscle cells to maintain vascular tone and healthy blood flow, we stimulated M_3_R, the sub-type predominantly expressed by SH-SY5Y cells(*33*), with a synthetic muscarine agonist, carbamoylcholine (carbachol), and measured HYZ-dependent changes in intracellular Ca^2+^ concentrations. Using Fura-2, a high-affinity ratiometric fluorescent indicator that specifically binds free Ca^2+^ ions inside cells, we showed that HYZ significantly reduced GPCR-dependent intracellular Ca^2+^ levels in a concentration-dependent manner, within the same concentration range that modulates RGS protein levels (Fig. 4E). Collectively, the results complement genetic data previously reported for ADO deficiency derived from the same cell line(*16*). Overall, the data explain the pharmacological activities of the drug and its molecular, cellular, and physiological mechanisms of action in evoking vasodilation for the treatment of preeclampsia.

Because G_αq_-coupled receptors mediate ERK phosphorylation(*34*) and are negatively regulated by RGS proteins(*35*), we next measured HYZ-dependent changes in ERK phosphorylation under the same conditions as above. Indeed, a concentration-dependent decrease in ERK phosphorylation was observed with HYZ in the presence of carbachol (Fig. 4F), suggesting that the side effect of drug-induced lupus may result from ADO inhibition and/or reduced ADO activity in patient T cells(*36–38*).

### ADO inhibition by HYZ senesces glioblastoma cells

Both elevated hypotaurine levels (the product of ADO’s metabolite substrate) and increased ADO expression have been clinically associated with glioblastoma growth and malignancy, with validated causal relationships(*39, 40*).This data suggests that ADO is a promising target for glioblastoma, however, no inhibitors for ADO have been reported to date. Given that HYZ is a > 70-year-old drug with a well-established safety profile, we explored ADO inhibition as a potential therapeutic strategy for glioblastoma first by treating glioblastoma cells lines with HYZ (Fig. 5A). A single treatment (10 and 100 μM) inhibited growth of U-87 cells throughout a time period of up to 9 days (Fig. 5B). Notably, chemoproteomics-based target identification experiments in U-87 cells, treated with the same concentration of HYZyne (100 μM), identified ADO as a high-occupancy target (fig. S17). At three days post-treatment, both U-87 and LN229 glioblastoma cell lines displayed concentration-dependent growth inhibition that plateaued at ∼ 30 μM with IC_50_ values ∼ 10 μM (Fig. 5C). The sensitivity of glioblastoma cell lines to HYZ was notably higher compared to non-cancer (HEK293T) and breast cancer cell lines (MDA-MB-231), which did not show a plateau and had estimated IC_50_ values of > 30 μM and > 50 μM for HEK293T and MDA-MB-231 cells, respectively. These results highlight HYZ’s cell-type selectivity toward glioblastoma cell lines.

**Fig. 5.**
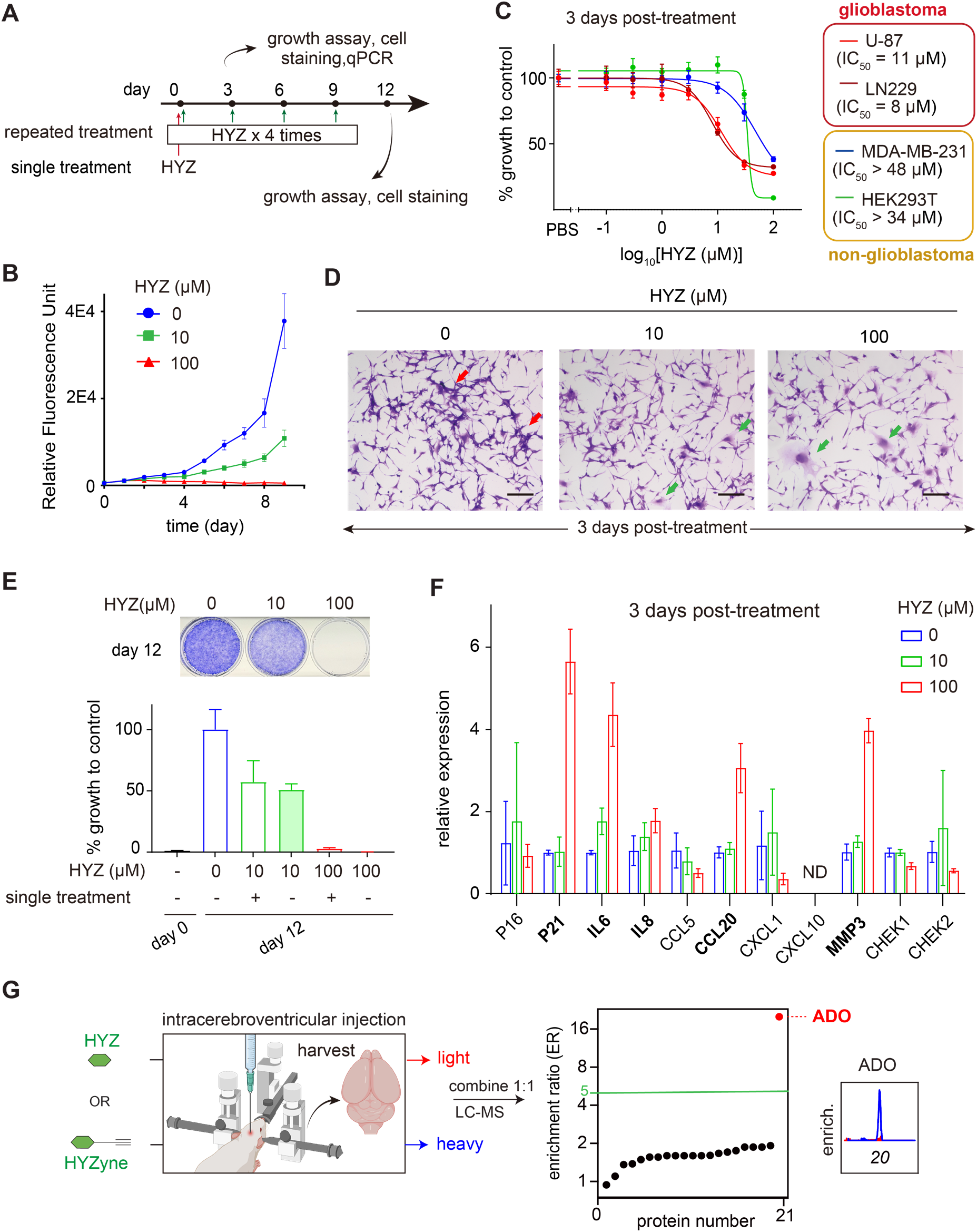
Glioblastoma growth inhibition by HYZ. (**A**) Schematic illustration of the experimental design of the study describing the HYZ treatment regimen and evaluation. (**B**) Quantification of U-87 cells over time from vehicle control and 10 or 100 μM HYZ treatment groups measured by alamarBlue assay. n = 5, mean ± SD. (**C**) Growth of varying cell lines 3 days post-treatment of HYZ. n = 4-5, mean ± SD. (**D**) Representative crystal violet staining of U-87 cells obtained from indicated HYZ treatment groups (0–100 μM). Red arrow: clusters of U-87 cells, green arrow: enlarged U-87 cells. Bar: 200 μm. (**E**) Growth of U-87 cells after 12-day culture with single and repeated HYZ treatments at 0–100 μM. Viable cells after repeated treatments are visualized by crystal violet staining (upper). Viable cells under various conditions were quantified by alamarBlue assay (lower). n = 6, mean ± SD. (**F**) qRT-PCR analysis of indicated senescence marker gene expression in U-87 cells after 3-day culture with 0–100 μM HYZ. n = 3, mean ± SD. (**G**) Quadrant plot of average enrichment ReDiMe ratios versus protein number for HYZyne from quantitative proteomics experiments in the soluble proteomes of mouse brain post intracerebroventricular injection of HYZ or HYZyne at 30 μg. Protein targets with ER ≥ 5 were considered high-reactivity targets and are annotated. [Part of figure created with BioRender.com]

Growth inhibition is represented by two general mechanisms: cytotoxic and cytostatic(*41*). We evaluated cytotoxicity at an early (6 h) time point, when cytotoxicity is typically observed(*42, 43*), and a later time point (24 h), when cell growth differences begin to emerge (Fig. 5B). No significant cell death was observed between treatment groups at either time point (fig. S18), suggesting HYZ is cytostatic rather than cytotoxic. Three days post-treatment, U-87 cells, which normally form cellular clusters, displayed morphological changes characterized by the absence of cluster formation and the presence of cellular enlargement with increasing HYZ concentrations (Fig. 5D). These features are often associated with cell-cycle arrest and cellular senescence. To investigate whether HYZ induces a senescent state, which is a common response to compounds with known cytostatic anti-cancer activity(*44*), we monitored cell growth over a 12-day (long-term) period following repeated treatments with HYZ (10 vs. 100 µM). No growth was observed at 100 µM (Fig. 5E). Because cellular senescence represents a terminal state of growth, we compared the cytostatic effects of single versus repeated treatments with HYZ. Notably, there was no substantial difference, further supporting the induction of senescence by a single treatment (Fig. 5E). Additionally, senescence marker p21, and senescence-associated secretory phenotype (SASP) markers including IL6, IL8, CCL20, and MMP3, were upregulated in a concentration-dependent manner after three days post-treatment (Fig. 5F), supporting that cellular senescence is the underlying mechanism of HYZ’s cytostatic effect.

### HYZ can inhibit ADO in the brain

HYZ itself has a poor blood-brain barrier (BBB) penetrance excluding delivery of the drug to the brain, thereby limiting its efficacy in treating the most severe and deadly symptoms of (pre)eclampsia (*e.g.,* seizure, coma, or cerebral hemorrhage) that can occur without warning. To ensure that HYZ can indeed engage ADO in the brain, we bypassed the BBB by delivering HYZyne/HYZ (30 μg) to mice by intracerebroventricular injection followed by global target analysis of brain tissue harvested from the sacrificed animals. The data show that ADO is the only detectable high-enrichment target from the brain proteome (Fig. 5G). As such, HYZ/HYZyne may be used as a pharmacological tool in the future to evaluate *in vivo* ADO activity in the brain and preclinical models of preeclampsia and/or glioblastoma.

## DISCUSSION

The results directly connect HYZ inhibition of ADO, increased accumulation of the known ADO substrates RGS4 and RGS5, attenuation of associated GPCR-mediated cell signaling, and subsequent reduction of intracellular Ca^2+^ levels, the key driver of vasodilation. This molecular, cellular and physiological mechanism for the pharmacological effect of HYZ complements previous reports of diminished RGS5 levels in the placenta and maternal arteries of preeclamptic mothers(*31*). Similarly, mice with the RGS5 gene disrupted exhibit the key characteristics of preeclampsia: hypertension, proteinuria, placental pathology, low birth weight, hyperresponsiveness to angiotensin II and glutamate signaling, blood brain barrier permeability and stroke(*31, 45*). Although comprehensive studies have not yet been performed, several lines of evidence also indicate that RGSs inhibit eclamptic seizures(*46, 47*), suggesting that a brain penetrant HYZ derivative could be effective against the most life-threatening complications of the disease, which include seizure, stroke and permanent organ damage(*48*).

The ability of this mechanism to explain existing genetic and (pre)clinical data reported for (pre)eclampsia is paralleled for glioblastoma but reversed with respect to what was known versus unknown. In preeclampsia, the drug was known and the target was unknown; whereas in glioblastoma, the target was known and the drug (or any drug leads) was unknown. As such, identifying the target in the former disease directed the discovery that HYZ arrested growth in the latter disease. Analogous to the case for preeclampsia, our biochemical data rationalize clinical data showing ADO expression and hypotaurine (the metabolic product of its activity) levels strongly correlate with tumor grade severity and patient outcome(*40, 49*), as also observed for other cancers(*50*).

In summary, by solving the > 70-year-old mystery around HYZ’s mechanism of action, we serendipitously identified the first inhibitor for an emerging therapeutic target for brain cancer. Matching the drug to its target provides a novel pharmacological tool to define the substrate scope of ADO and elucidate its function in other diseases. It will also guide the improvement of its properties and provide a tool to solve the 2400-year-old mystery of (pre)eclamptic etiology.

## Acknowledgements

We thank C. Brisson and T. Ibrahim for computational support (School of Arts and Science Computing, University of Pennsylvania), L. He (Microsoft), R. Park (Integrated Proteomics Applications, Inc.) and R. Suciu (Scripps Research) for proteomics data analysis tools, P. E. Dawson (Scripps Research) for assistance with the preparation of the TEV tags, K. S. Zarret and K. Ito (University of Pennsylvania) for providing access to an Eclipse TE2000-U for cell imaging and M. C. Simon (University of Pennsylvania) for generous support and mentorship in designing senescence assays.

## Author contributions

K.S. and M.L.M. designed experiments and interpreted results. X.W., N.R.M. and R.H. synthesized probe. K.S., K.A.B., E.W.B., S.R.C., M.A., S.W.K, Z.L., R.H. and N.R.M. performed gel- and MS-based experiments. K.S., K.A.B. and M.L.M. analyzed MS data. J.L. and A.L. performed inhibition assay with isolated human ADO, crystallography, and EPR studies. Y.C. and K.A.M. performed and analyzed GPCR signaling assays, K.H. and K.S. performed intracerebroventricular drug delivery experiments, K.S. and N.R.M. performed RGS and calcium signaling assays and growth inhibition studies. K.S. and E.W.B. performed and analyzed senescence data. K.S. and M.L.M. wrote the manuscript. K.S., M.L.M, J.L., A.L., Y.C., K.A.M., Z.A.B., J.M.B., W.H.P., E.W.B and D.M.O. edited the manuscript and all authors reviewed it.

## Data and materials availability

PDB files have been deposited into RCSB with the accession number 9DMA and 9DY4, respectively. All other data are available in the main text or the supplementary materials.

## Supplementary Materials

Materials and Methods

Figs. S1 to S18

Tables S1 to S3

Data S1 to S3

