## Supplemental material for "Hydralazine inhibits cysteamine dioxygenase to treat preeclampsia and senesce glioblastoma"

**The PDF file includes:**

Materials and Methods
Figs. S1 to S18
Tables S1 to S3
References

**Other Supplementary Materials for this manuscript include the following:**

Data S1 to S3

### Materials and Methods

#### Clickable probes and drugs

Clickable chemical probes were synthesized and characterized according to previous methods(*11*). All probes and drugs (HYZ and HYZyne) were prepared fresh from solid stocks in PBS and adjusted to pH ~7 with 5 M NaOH the day of experiments. HYZ and HYZyne are soluble and stable at 50–100 mM in PBS at pH~7.

#### Cell lines and culture conditions

All cells in this study were purchased from American Type Culture Collection (ATCC). HEK293T, MDA-MB-231, U-87, LN229, RAW 264.7 and Neuro 2a cells were grown at 37 °C in a humidified 5% $CO_2$ atmosphere and expanded in high-glucose DMEM supplemented with 10% (v/v) fetal bovine serum (FBS). SH-SY5Y cells were cultured in DMEM/F12 medium supplemented with 10% FBS. For Stable Isotope Labeling by Amino acids in Cell culture (SILAC) experiments, each cell line was passaged a minimum of six times in lysine- and arginine-free DMEM (Thermo Fisher Scientific #A33822) containing dialyzed FBS (Silantes #281001200), and isotopically enriched *L*-[$^{13}C_6{}^{15}N_2$]lysine hydrochloride and *L*-[$^{13}C_6{}^{15}N_4$]arginine hydrochloride (100 µg/mL each, Sigma) or natural abundance isotopologues (100 µg/mL each, 550 µM and 475 µM, respectively, Sigma) as described previously(*10*). All cells were routinely tested for the absence of mycoplasma contamination with MycoStrip (Invitrogen, REP-MYS-20).

#### *in situ* labeling of cells to identify HYZ-reactive targets

*in situ* labeling of cells for gel- and MS-based experiments was conducted following previously described protocols with some exceptions(*10, 14*). For gel-based experiments with the endogenous proteome, cells were treated with either HYZ or HYZyne at a concentration of 10 µM for 30 minutes instead of 1 mM. To assess the effect of iron chelator and site-directed mutagenesis, cells were treated with HYZyne at 100 µM unless otherwise noted. For MS-based experiments with U-87 cells, 100 µM HYZyne and HYZ were used for enrichment instead of 1 mM, while 100 µM HYZyne and 500 µM HYZ were used for the competition experiments instead of 1 mM and 10 mM, respectively.

Cell proteomes for gel- and MS-based experiments were prepared as described previously(*10*). For SILAC experiments, isotopically heavy and light whole cell lysates were mixed in equal proportions prior to fractionation of membrane and cytosolic (soluble) components.

### Gel-based analysis of HYZyne-labelled proteins

To analyze protein activity, soluble and membrane tissue or cell proteomes were conjugated to rhodamine-azide by copper-catalyzed azide-alkyne cycloaddition (CuAAC or 'click') chemistry, resolved by SDS-PAGE and visualized by in-gel fluorescence scanning as previously described (*14*), with the modification of using 2.5 µM azide-rhodamine (0.1 µL/sample, 1.25 mM in DMSO) for endogenous proteome instead of the originally described 25 µM.

### MS-based quantitative analysis of HYZyne-labelled proteins

Proteomes were similarly conjugated to biotin-azide by CuAAC, probe-labelled proteins were enriched by streptavidin affinity chromatography and digested on-bead with trypsin as previously described(*10, 14*). For mouse tissue proteome, the resulting tryptic peptide digests from probe- versus drug-treated mice were isotopically differentiated by reductive dimethylation (ReDiMe) of lysine residues using isotopically 'heavy' and 'light' formaldehyde, respectively to identify and quantify enrichment heavy/light ReDiMe ratios for HYZyne as previously described (*13*).

### Animals for target identification *in vivo*

C57BL/6NJ mice (8–14 weeks) were purchased from Jackson Laboratories and used for *in vivo* experiments. All mice were housed with 2–5 mice per cage under a 12-hour normal light-dark cycle (light on AM, light off PM) with ad-libitum access to standard chow and water. Protocols were approved by the Institutional Animal Care and Use Committees (IACUC) at the University of Pennsylvania (Philadelphia, PA).

### Identification of HYZyne-enriched targets from heart tissue

For target identification in heart proteome, C57BL/6NJ male mice were treated intraperitoneally with the HYZ or HYZyne (12.5 mg/mL) for 4 h at 50 mg/kg, following previously described proteome preparation protocols with slight modifications(*13*). Modification includes the use of zirconium oxide beads (2 mm, Next Advance, Inc.) for homogenization.

### Liquid chromatography-tandem mass spectrometry (LC-MS/MS) analysis of tryptic digests

Peptide mixtures from tissue or cell proteomes were desalted prior to identification (based on MS2 spectra assignments) and quantification (based on intensity of corresponding extracted MS1 parent ion chromatograms) by liquid chromatography-tandem mass spectrometry-based proteomics (LC-MS/MS), and analyzed according to previously reported methods(*14, 51, 52*). First, proteomes analyzed following treatment of the probe (defined by its alkyne handle for biochemical enrichment) must ‘enrich’ for the protein of interest when compared against the native drug (which has no handle for enrichment). If sufficient enrichment is achieved from cells harvested from probe-treated cells *versus* that of native drug-treated cells (enrichment ratio or ER), the protein can then be evaluated for ‘competition’ by pretreating cells with the drug prior to treatment with the probe. If the drug blocks enrichment by the probe (competition ratio or CR), this criterion indicates high reactivity of the target towards the drug and high-occupancy of the target by the drug. This parameter is critical for determining small molecule-target engagement and stoichiometry and is useful in assessing the likelihood that engagement of a specific target may produce a therapeutic response (*e.g.,* efficacy) or cause deleterious side effects in patients (*e.g.,* safety). Proteins both substantially enriched (MS1median heavy: light SILAC or ReDiMe ER > 5) and competed by the drug (MS1 median heavy: light SILAC or ReDiMe CR > 3) were considered high-reactivity and high-stoichiometry targets of HYZyne. Median heavy/light SILAC or ReDiMe peptide ratios were derived from three or more unique quantified peptides per protein and averaged across replicates to generate the final ratios(*13*). For mouse tissues, protein ER and CR values were determined using the median ratio from two or more unique peptides, while for U-87 cells, one or more unique peptides were used.

**Cloning and transfection of overexpression constructs in HEK29T cells**

The full-length human gene encoding ADO was amplified from a cDNA library derived from the same low-passage HEK293T cells used for proteomic profiling as reported previously(*14*). The coding sequence cloned was matched to ADO coding sequence in GenBank (BC018660.1 and AK127694.1). The sequence has two base pair mutations yielding Gly25Trp and Pro39Ala mutants compared to NCBI reference ADO sequence (NM_032804.6). Human RGS5 cDNA was purchased as the pcDNA3.1+c-(k)-DYK (GenScript, OHu22885) and subcloned into our pRK5-flag vector (N or C-terminal FLAG tag) for mammalian expression (*14*). RGS4/16 Active-site mutants of ADO were generated using Quikchange XLII site-directed mutagenesis (Agilent) using primers (Integrated DNA Technologies) containing the desired mutations. All constructs were verified by DNA sequencing (Penn Genomic and Sequencing Core). Low-passage HEK293T cells were grown to 40~60% confluency under standard conditions. ADO and RGS4/5 were transiently overexpressed with polyethyleneimine (PEI) 'MAX' (MW 40,000, Polysciences Inc.) as previously described(*14*). All proteins were overexpressed in HEK293T cells contained an addition 15 amino acids appended to their *C*-terminus ($A_3G_4DYKD_4K$).

**Site- and structure-based resolution of HYZyne-captured peptide from ADO-overexpressing cells**

To identify the basis for ADO inhibition by HYZyne and HYZ, we isolated and characterized the HYZyne-modified peptides from probe-treated HEK293T cells overexpressing recombinant human ADO as adopted from previous studies (*14, 19*). Briefly, the resultant probe-labelled peptides are recognized as co-eluting isotopic pairs in a 1:1 ratio that are resolved by 6.0138 Da due to incorporation of natural abundance (light) or L-[$^{13}C_5$ $^{15}N$]valine (heavy) into the portion of the tag that is retained, enabling unbiased labeled peptide identification without prior knowledge.

**Purification and reconstitution of active ADO from *E. coli***

Bacterial plasmids with a TEV protease cleavage site containing WT human ADO (hADO) for assays and spectroscopic studies and C18S/C239S hADO variant for crystallizations have been described elsewhere(*23, 24*). The plasmids were transformed into BL21 (DE3) (Merck) for cell culture. Cell cultures were prepared at 37 °C in an M9 minimal medium containing Kanamycin (50 µg/ml), with shaking (220 rpm), until an optical density of 0.8 AU at 600 nm was reached. Following 5 h induction with 20 µM $CoCl_2$ or $(NH_4)_4Fe(CN)_6$ and 0.5 mM isopropyl-β-thiogalactoside (IPTG) at 28 °C, the cells were harvested and resuspended in lysis buffer (50 mM Tris-HCl and 200 mM NaCl at pH 8.0) and then disrupted by a Microfluidizer LM20 cell disruptor. The supernatant was recovered after centrifugation (34,000×g for 40 min) at 4 °C. The Co-NTA agarose beads were used for Co(II)•ADO purification, while he Ni-NTA agarose beads were used for Fe(II)•ADO purification. After washing the column with washing buffers (50 mM Tris-HCl, 200 mM NaCl, and 10 mM imidazole, pH 8.0), the protein was eluted with elution buffer (50 mM Tris-HCl, 200 mM NaCl, and 300 mM imidazole, pH 8.0). The *N*-terminal $His_6$-tagged ADO was mixed with the appropriate TEV protease and dialyzed in a buffer containing 10 mM Tris-HCl and 5% glycerol (pH 8.0) overnight. After removing the $His_6$-tag and uncut protein by the IMAC column, the non-tagged hADO proteins were further purified using in-house Superdex 75 (Cytiva) gel-filtration column (16/600, 1200 mL), equilibrated with a buffer containing 50 mM Tris-HCl and 50 mM NaCl (pH 7.6). After verification by SDS-PAGE, the purified proteins were either concentrated to the required concentration for subsequent experiments or stored in a buffer containing 50 mM Tris-HCl, 50 mM NaCl, and 5% glycerol (pH 7.6) at –80 °C. The protein concentration was determined based on the extinction coefficient of $\varepsilon_{280\ nm}$ = 22,920 $cm^{-1}M^{-1}$.

**Structural characterization of HYZ•Co(II)• or Fe(II)•ADO complex by X-ray crystallography**

The crystallization was achieved using the hanging drop vapor-diffusion method at 289 K with a buffer continuing 0.1 M BisTris-HCl (pH 5.5), 0.2 M $(NH_4)_2SO_4$, and 20% (w/v) PEG3350, Microcrystals formed after 1–2 days and reached an optimal size for X-ray diffraction after seven days. The Co(II)•ADO or Fe(II)•ADO crystals were soaked with HYZ (30 min, 50 mM), then cryoprotected with crystallization buffer containing an additional 25% (v/v) glycerol before being flash-cooled in liquid $N_2$. Crystallographic data were collected at 100 K temperature at Stanford Synchrotron Radiation Lightsource (SSRL) beamline BL 9-2. All X-ray diffraction intensity data were integrated, scaled, and merged using HKL3000(*53*). Molecular replacement was performed with Phenix(*54*) using the structure of Ni-hADO (PDB entry 7REI)(*55*) as a starting model. The final model was manually adjusted and further refined with Coot(*56*) and Phenix(*54*). Ramachandran statistics were analyzed using MolProbity (http://molprobity.biochem.duke.edu/). All the molecular model figures were generated using PyMOL (W.L. DeLano, The PyMOL Molecular Graphics System version 1.8.6.0. Schrödinger LLC, http://www.pymol.org/; 2002). The structural data was deposited to the PDB databank with entry codes 9DMA and 9DY4.

#### Characterization of HYZ binding to Co(II)•ADO by EPR spectroscopy

A stock solution of HYZ (100 mM) or cysteamine (100 mM) was prepared anaerobically and added to the Co(II)•ADO to reach the final concentration under anaerobic conditions. The samples were transferred to quartz EPR tubes and slowly frozen in liquid nitrogen. EPR spectra were recorded on a Bruker E560 X-band spectrometer equipped with a cryogen-free 4 K temperature system with an SHQE high-Q resonator, operating at 100 kHz modulation frequency, with 0.6 mT modulation amplitude, and averaging four scans per spectrum. Full-range scans were conducted at 3.17 mW microwave power at 30 K.

#### Inhibition of the dioxygenase activity of isolated Fe(II)•ADO activity by HYZ

The inhibition assays were conducted using an oxygen electrode (Oxygraph, Hansatech Instruments). The iron content of the enzymes was determined using the Ferrozine assay(*57*). The enzyme (2 µM) was pre-incubated with excess ascorbic acid (20 µM) and HYZ (0, 6.7, or 10 µM). Standard conditions were used with enzyme (2 µM), Tris-borate buffer (100 mM), and varying concentrations of cysteamine (0–48 mM) in a total volume of 0.5 mL (pH 8.0, 25 °C). The reaction was initiated by adding 10 µL cysteamine to the electrode chamber while maintaining a constant stirring speed of 100 rpm. Net oxygen consumption was determined by subtracting the initial oxygen consumption rate in the absence of the enzyme from that in the presence of the enzyme. All experiments were performed in triplicate. The three lines were generated by plotting the reciprocal of net oxygen consumption against the reciprocal of enzyme concentration at different HYZ concentrations. Global fittings were conducted across the datasets using OriginPro (OriginLab), where the parameters $k_{cat}$, $K_M$, $K_{i1}$, and $K_{i2}$ are shared in the following equation:

$$\frac{1}{v_0} = \left[\left(1 + \frac{[I]}{K_{i1}}\right)\frac{K_m}{k_{cat}}\frac{1}{[S]}\right] + \left(1 + \frac{[I]}{K_{i2}}\right)\frac{1}{k_{cat}}$$

#### Harvesting cells and expression analysis by Western Blotting

After treatment with various concentrations of HYZ, HEK293T and SH-SY5Y cells in 6 cm dishes were washed with PBS (3 × 3 mL). Then, cells were collected by scraping using RIPA lysis and extraction buffer (300 µL, Thermo #89900). Lysates were then sonicated to shear genomic DNA, and the protein concentration was quantified by DC protein assay kit (Bio-Rad #5000111) on ELx808 plate reader (Biotek).

Protein expression was analyzed as previously described(*10*). The primary antibodies and dilutions were as follows: anti-FLAG (1:2500, F1804, Sigma), anti-ADO (1:1000, ab134102, Abcam), anti-RGS4 (1:250, D4V1P, Cell Signaling), anti-RGS5 (1:100, sc-514184, Santa Cruz), anti-RGS16 (1:100, sc-166083, Santa Cruz), anti-ERK1/2 (1:2000, 9102, Cell Signaling), anti-phospho-ERK1/2 (1:2000, 9101, Cell Signaling), anti-GAPDH (1:1000, MA515738, Invitrogen). The secondary antibodies and dilutions were as follows: goat anti-mouse HRP (1:10000, ab150113, Abcam), goat anti-rabbit HRP (1:2000, 65-6120, Invitrogen), goat anti-rabbit Alexa Fluo 488 (1:2000, A-11034, Invitrogen), sheep anti-mouse HRP (1:2000, NA931V, Sigma).

#### Determination of $IC_{50}$ values

After *in vitro* competitive labeling by SDS-PAGE gel and visualization of labeled proteins via in-gel fluorescence scanning, the percentage of remaining activity at active site was determined by measuring the integrated optical intensity of the bands as described previously(*11*). ADO activity inhibition by HYZ was similarly assessed by measuring RGS protein levels through Western blot, quantified by the integrated optical intensity of the bands. Nonlinear regression (Fit curve) analysis was used to determine the $IC_{50}$ values, which was derived from a dose-response inhibition curve generated using GraphPad Prism 10.

#### Cell-based GAP assay

The overexpression plasmids used in GPCR signaling (BRET) assay were generated as previously described(*32*), and combined with our *C*-terminally flagged tagged ADO expression vector described above. The impact of HYZ on GAP activity was measured via the BRET assay, performed as previously described(*32*). Briefly, HEK293T/17 cells ($2 \times 10^6$ per well) were plated in 0.1% Matrigel (Corning #356230) coated 35 mm plates in 1.5 ml of growth media and transfected for 4 h. Constructs were transfected using PLUS (5 µl) and Lipofectamine LTX (6 µl) (Invitrogen #15338-100) per plate as following: $M_3R$ (0.21 µg), Venus 156-239-G$\beta_1$ (0.21 µg), Venus 1-155- G$\gamma_2$ (0.21 µg), masGRK3ct-Nluc-HA (0.21 µg), G$\alpha_q$ (0.42 µg), RGS4 (2.52 µg), RGS5 (2.52 µg), RGS16 (0.63 µg), and ADO (1.22 µg). pcDNA3.1+ was used to normalize the amount to 5 µg for each transfection. 24 h after transfection, cells were washed once with BRET buffer (PBS containing 0.5 mM $MgCl_2$ and 0.1% glucose) and collected by pipetting. Harvested cells were washed by centrifugation (500 g, 5 min, 4 °C) and resuspended in BRET buffer. Cells ($5 \times 10^4$ per well) were plated in 96-well flatbottomed white microplates (Greiner Bio-One) and cells were incubated with buffer or various concentration of HYZ for 1 h. After addition of the NanoLuc substrate (Promega #N1120), BRET measurements were made using a microplate reader (POLARstar Omega; BMG Labtech). Acetylcholine (100 µM) and atropine (100 µM) were injected sequentially for activation and inhibition $M_3R$, respectively. The BRET signal is calculated as the ratio of the light emitted by the acceptor (535 nm with a 30 nm band path width) over the light emitted by the donor (475 nm with a 30 nm band path width). BRET ratio prior to the injection of agonist was used for baseline correction and the resulting ΔBRET ratio was normalized to maximal ΔBRET reading upon agonist stimulation. The rate constants ($1/\tau$) of the deactivation phases were obtained by fitting the one phase decay model to the traces using GraphPad Prism 10.

#### Intracellular calcium measurement

SH-SY5Y cells ($7 \times 10^4$) were plated and cultured in 96-well plate (Sigma-Aldrich #CLS3603) for 24 h at 37 °C with 5% $CO_2$ in DMEM/F12 supplemented with 10% FBS (100 µL). Cells were treated with reagents of Fura-2 assay kit (abcam #ab176766) and HYZ at various concentrations (0-100 µM) according to the manufacturer's instructions (1 h). Following incubation, intracellular $Ca^{2+}$ levels were measured by detecting the fluorescence intensity at 510 nm, excited alternately at 340 nm ($F_{340}$) and 380 nm ($F_{380}$) every 21 s with the Infinite M1000 (TECAN) at 37 °C. Total $Ca^{2+}$ measurement time frame was from 0 to 532 s, whereas basal $Ca^{2+}$ concentration was measured between 0 to 128 s. Carbachol (100 µM) was added to induce GPCR-dependent $Ca^{2+}$ signaling at 128 s. $F_{340}/F_{380}$ ratios were quantified to represent intracellular $Ca^{2+}$ levels. To determine the maximum response to carbachol, the minimum $F_{340}/F_{380}$ ratios ($F_{min}$) during the basal period (0–128 s) was subtracted from the maximum $F_{340}/F_{380}$ ratios ($F_{max}$) observed after carbachol addition (149–532 s).

#### Cell growth and proliferation assays

For the 3-day culture, cells (1500) were seeded in 96-well plates (Sigma-Aldrich #CLS3603) and incubated at 37 °C with 5% $CO_2$ in DMEM (100 μL) supplemented with 10% FBS for 24 h. Cells were treated with various concentrations of HYZ or PBS and incubated for 3 days. Cell growth was quantified using the alamarBlue assay as described below. For morphological and gene expression analysis, cells ($2 \times 10^5$) were plated in 6 cm dishes in the media (3 mL), cultured for 3 days under the same conditions, and analyzed by crystal violet staining or qPCR.

For the 9-day time course experiment, U-87 cells (500) were seeded in 96-well plates and incubated under the same conditions for 24 h. Cells were treated with various concentrations of HYZ and incubated for 0–9 days. Each day, 5 samples per group were assessed using the alamarBlue assay. Cells used for one time point were not reused on subsequent days, and the media was not refreshed during the time course.

For the 12-day (long-term) culture, U-87 cells (250 for 96-well plates or $1 \times 10^4$ for 6 cm dishes) were seeded and cultured in DMEM with 10% FBS (100 μL for 96-well plates or 3 mL for 6 cm dishes) for 24 h. Cells were treated with various concentrations of HYZ or PBS and incubated for 12 days, with media and HYZ refreshed every 3 days unless otherwise noted. Cell growth in 96-well plates was assessed using the alamarBlue assay, while cells in 6 cm dishes were subjected to crystal violet staining.

#### alamarBlue assay

Cell viability was quantified based on NADH levels in live cells. Cells in 96 well plates (100 μL media) were treated with alamarBlue HS (3 h, 10 μL) (Invitrogen #A50100). Fluorescent signals were measured using an excitation of 560 nm and emission of 590 nm with the Infinite M1000 (TECAN). Background fluorescence from media, averaged from five wells containing media (100 μL) and alamarBlue HS (10 μL) without cells, was subtracted from the fluorescence intensity values of each sample.

#### Crystal violet staining

Cell viability and morphology were evaluated with crystal violet, which stains both pan DNA and proteins. Cultured cells were fixed with 4% paraformaldehyde (PFA) for 10 min and stained with crystal violet solution [0.5%, methanol:water = 20:80 (v/v)] (Sigma-Aldrich #C6158) for 30 min at room temperature. Cells were then washed with tap water to remove residual crystal violet. For morphological study, images were acquired by Eclipse TE2000-U (Nikon).

#### Measurement of cell death

Cytotoxicity was quantified using double staining with live-cell permeable and impermeable DNA dyes. U-87 cells were plated and cultured for 24 h at 37 °C with 5% $CO_2$. Cells were treated with HYZ (0–100 μM) for 6 or 24 h. Cells were stained with Hoechst33342 (10 μg/mL) (BD Pharmingen #BDB561908) and PI (1 μg/mL) (BD Pharmingen #BDB556463) for 30 min at 37 °C. Positive control cells were simultaneously treated with 0.1% Triton X-100. Images were acquired by Eclipse TE2000-U (Nikon).

### qPCR

mRNA expression levels were measured using real-time quantitative RT-PCR (qPCR). Total RNA was purified from cells with the RNeasy Mini Kit and DNase I (Qiagen #74104 and 79254). cDNA was synthesized from 1 μg mRNA with iScript™ Reverse Transcriptase (Bio Rad #1708841). qPCR was performed with QuantStudio 6 Flex Real-Time PCR System (Thermo) and Luna Universal qPCR Master Mix (NEB #M3003). Cycle threshold (Ct) values for senescence markers expression in U-87 cells were normalized to Ct values of 18S ribosomal RNA. Relative expression levels of each marker were calculated using the ΔΔCt method and normalized to the average values of the vehicle controls. Markers with undetermined Ct values due to high cycle numbers were classified as “not detected” in the figure. Primer sequences were based on a previous literature(*58*) and provided in Data S3.

### Target identification in the brain by intracerebroventricular delivery of HYZyne vs. HYZ

For the intracerebroventricular brain delivery experiment only, NSG (NOD scid gamma mouse) mice (6–14 weeks) were obtained from the Stem Cell and Xenograft Core at the University of Pennsylvania. Anesthesia was induced with inhaled isoflurane (1–4%) and confirmed via pedal reflex before incision. The surgical area was prepared by shaving the head (Wahl Arco Cordless Clipper) and sterilizing the skin with alcohol prep pads. A midline incision was made using a #10 scalpel. A digital stereotaxic frame (Stoelting #51730D) and hand drill (Stoelting #51449) were used to create burr holes in the skull. The bregma was identified, and a 10 μL Hamilton syringe (Hamilton #80301) was positioned 1 mm lateral to the left and 0.3 mm anterior to bregma. The syringe was lowered until it touched the brain surface, zeroed, and advanced 3 mm deep. HYZyne or HYZ was injected at 1.5 μL (30 μg dissolved in 0.9% saline, adjusted to pH ~7 with 5M NaOH). The incision was closed with skin glue (3M Animal Health #1469SB), and mice were allowed to recover. After 4 h, mice were euthanized, brain tissues were harvested and proteomes were processed, ratiometrically compared and analyzed as described previously (*13*).

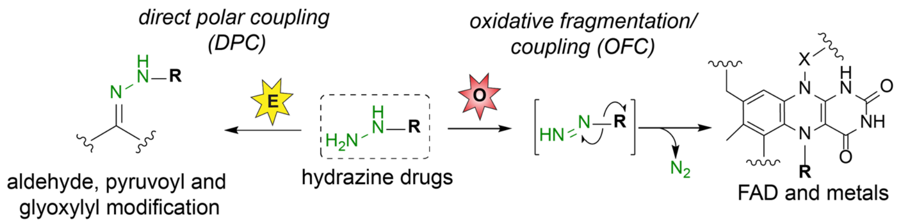

**Fig. S1. Modes of hydrazine reactivity.** Schematic for two distinct modes of probe reactivity: direct polar coupling (DPC) and oxidative fragmentation/coupling (OFC). Green depicts the reactive probe moiety.

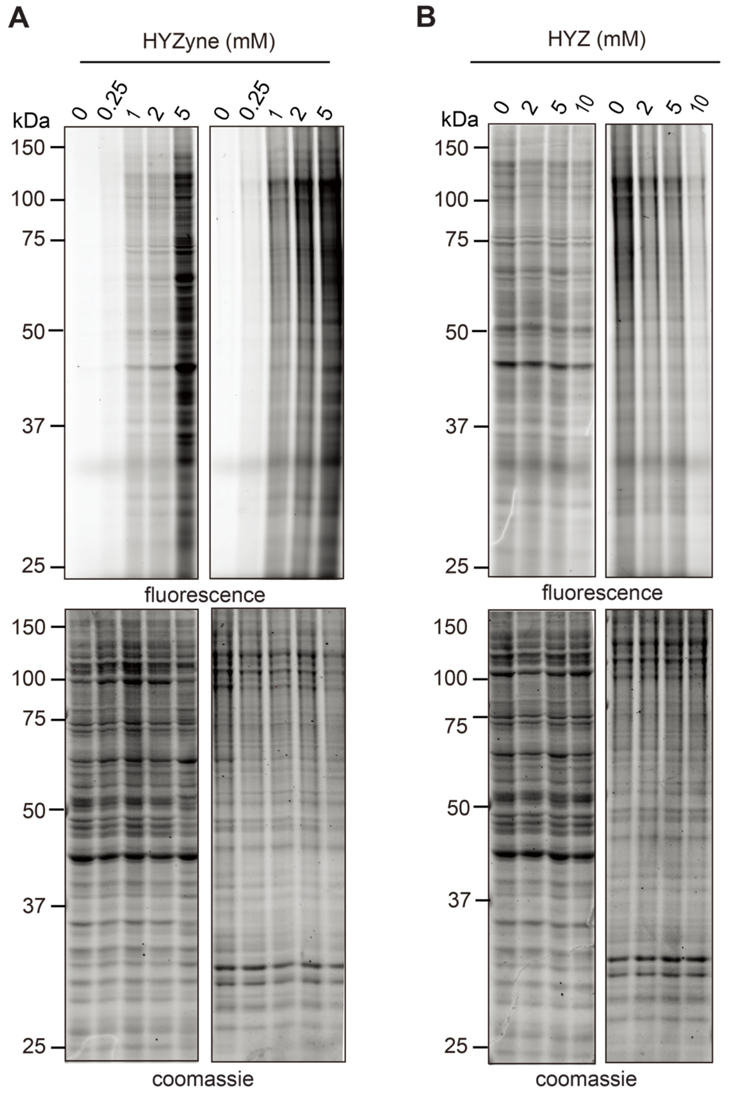

**Fig. S2. Gel-based profiles for HYZyne-treated HEK293T cells.** (**A**) Concentration-dependent labelling profiles for soluble (left) and membrane proteomes (right) of HYZyne-treated HEK293T cells (upper). Corresponding expression profiles are shown (lower). (**B**) Competition of HYZyne labelling by HYZ for soluble (left) and membrane proteomes (right) of HEK293T cells co-treated with 1 mM HYZyne and varying concentrations of HYZ (upper). Corresponding expression profiles are shown (lower).

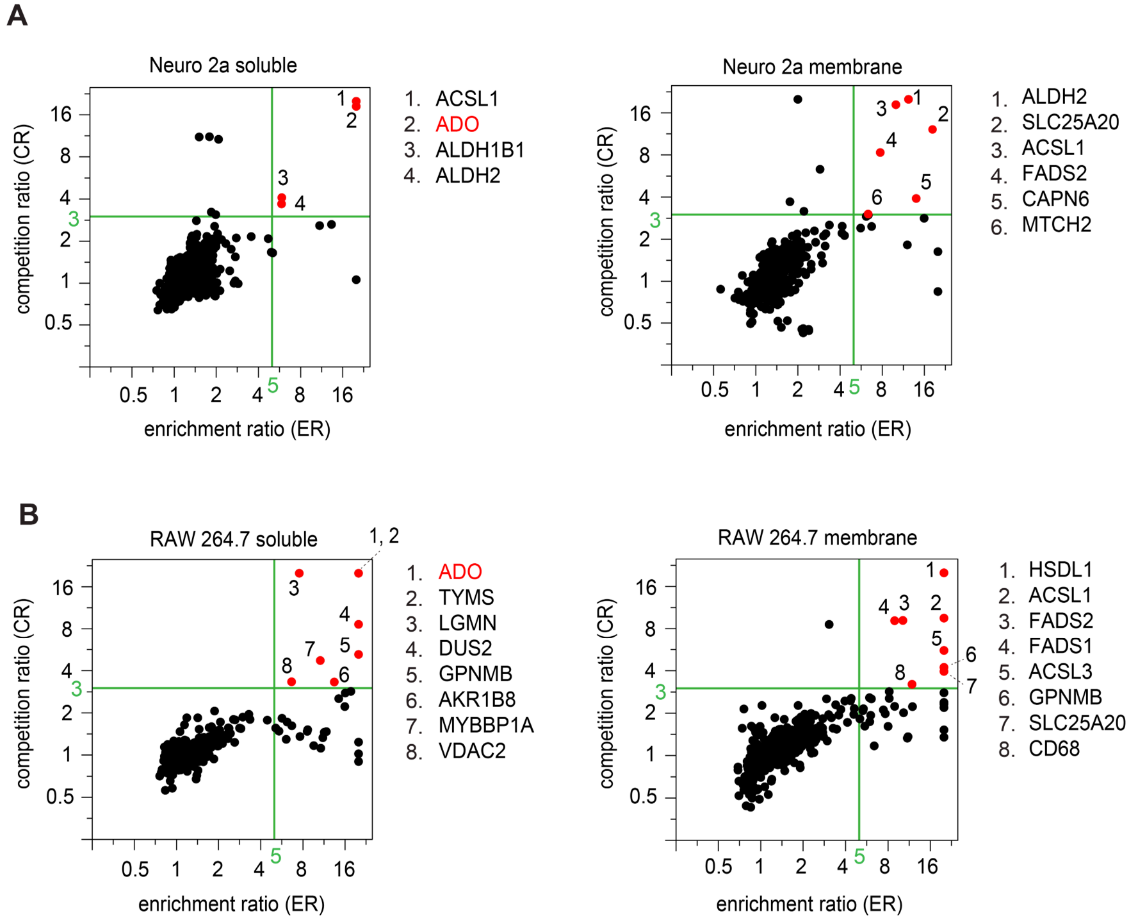

**Fig. S3. Identification of protein targets of HYZyne in two mouse cell lines.** (**A, B**) Quadrant plot of average competition versus average enrichment SILAC ratios from quantitative proteomics experiments for the soluble (left) and membrane (right) proteomes of Neuro 2a (**A**) or RAW264.7(**B**) cells. Proteins with ER ≥ 5 and CR ≥ 3 (upper right quadrant) were considered high-occupancy targets; listed to the right of the plot. Proteins highlighted red are investigated in this work.

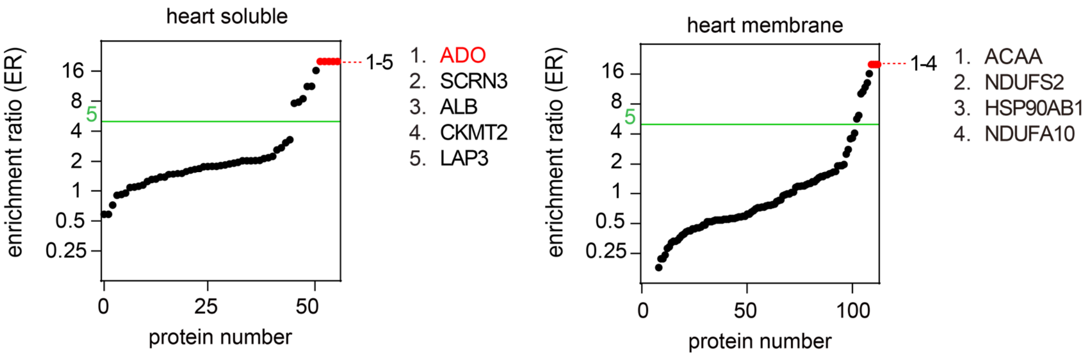

**Fig. S4. High-reactivity targets of HYZyne in mouse heart.** Average enrichment ReDiMe protein ratios versus protein number for HYZyne from quantitative proteomics experiments in the soluble (left) and membrane (right) proteomes of mouse heart 4 h post-injection via ratiometric comparison of HYZ (light) versus HYZyne (heavy) (50 mg/kg, intraperitoneally). Protein targets with ER ≥ 5 were considered high-reactivity targets. Proteins with ER ≥ 20 are annotated and are listed on the right of the plot. Proteins highlighted red are investigated in this work.

A

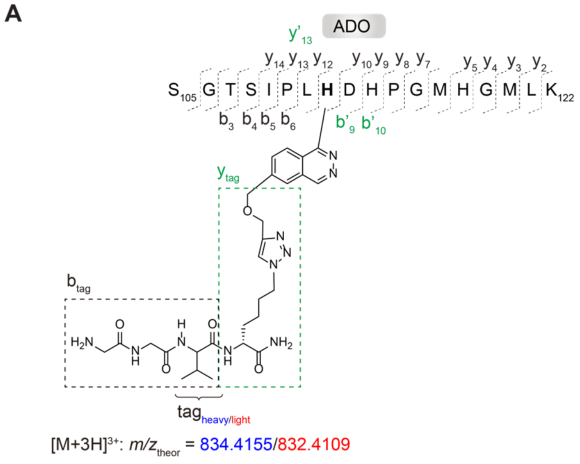

B

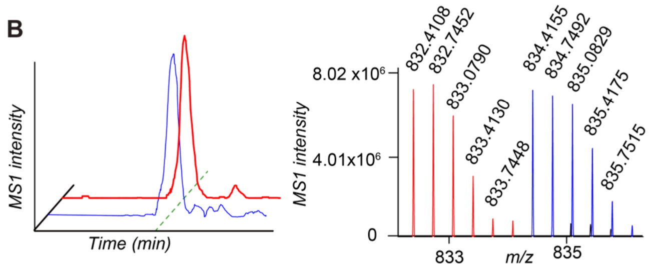

C

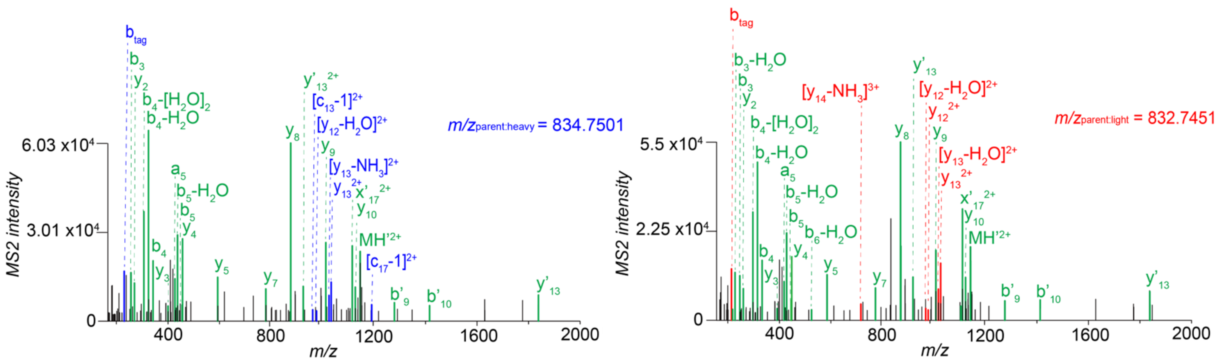

D

| | Theoretical $m/z$ Heavy | Theoretical $m/z$ Light | Measured $m/z$ Heavy | Measured $m/z$ Light |
|---|---|---|---|---|
| $b_{tag}$ | 220.1330 | 214.1192 | 220.1332 | 214.1188 |
| $[c_{13}-1]^{2+}$ | 967.4777 | 964.4708 | 967.4709 | - |
| $[c_{17}-1]^{2+}$ | 1186.5802 | 1183.5733 | 1186.5763 | - |
| $y_{13}^{2+}$ | 1029.0150 | 1026.0081 | 1029.0092 | 1026.0079 |
| $[y_{13}-H_2O]^{2+}$ | 1020.0097 | 1017.0028 | 1019.9988 | 1016.9901 |
| $[y_{13}-NH_3]^{2+}$ | 1020.5017 | 1017.4948 | 1020.5014 | 1017.4931 |
| $y_{12}^{2+}$ | 980.4886 | 977.4817 | - | 977.4818 |
| $[y_{12}-H_2O]^{2+}$ | 971.4833 | 968.4764 | 971.4791 | 968.4714 |
| $[y_{14}-NH_3]^{3+}$ | 718.3649 | 716.3603 | - | 716.3618 |
| $a_3$ | 218.1135 | | - | 218.1133 |
| $a_5$ | 418.2296 | | 418.2299 | 418.2303 |
| $b_3$ | 246.1084 | | 246.1086 | 246.1086 |
| $b_3$-$H_2O$ | 228.0979 | | - | 228.0980 |
| $b_4$ | 333.1405 | | 333.1407 | 333.1405 |
| $b_4$-$H_2O$ | 315.1299 | | 315.1299 | 315.1302 |
| $b_4$-$[H_2O]_2$ | 297.1196 | | 297.1194 | 297.1196 |
| $b_5$ | 446.2245 | | 446.2246 | 446.2239 |
| $b_5$-$H_2O$ | 428.2140 | | 428.2141 | 428.2147 |
| $b_6$-$H_2O$ | 525.2667 | | - | 525.2672 |
| $b'_9$ | 1275.6229 | | 1275.6292 | 1275.6240 |
| $b'_{10}$ | 1412.6818 | | 1412.6884 | 1412.6901 |
| $y'_{13}$ | 1836.8897 | | 1836.8807 | 1836.8845 |
| $y_{10}$ | 1122.5183 | | 1122.5189 | 1122.5204 |
| $y_{10}$-$H_2O$ | 1104.5077 | | - | 1104.5116 |
| $y_9$ | 1007.4914 | | 1007.4906 | 1007.4922 |
| $y_8$ | 870.4324 | | 870.4326 | 870.4330 |
| $y_7$ | 773.3797 | | 773.3797 | 773.3790 |
| $y_5$ | 585.3177 | | 585.3179 | 585.3179 |
| $y_4$ | 448.2588 | | 448.2591 | 448.2590 |
| $y_3$ | 391.2374 | | 391.2366 | 391.2362 |
| $y_2$ | 260.1969 | | 260.1969 | 260.1971 |
| $MH'^{2+}$ | 1141.5571 | | 1141.5586 | 1141.5574 |
| $y'^{2+}_{13}$ | 918.9485 | | 918.9494 | 918.9517 |
| $y_8^{2+}$ | 435.7199 | | 435.7203 | 435.7186 |

**Fig. S5. MS/MS characterization of HYZyne-labelled active-site peptide of ADO.** (**A**) Structures and theoretical parent masses of heavy- and light-tagged ADO peptides labelled by HYZyne and processed by the isoTOP-ABPP method. (**B**) **E**xtracted parent ion chromatograms and corresponding isotopic envelopes for heavy- (blue) and light- (red) tagged peptides quantified in human ADO-transfected HEK293T cells. (**C**) MS2 spectra generated from indicated parent ions (left: heavy and right: light). Unshifted ions are shown in green whereas fragment ions that retain the heavy or light portions of the tag are shown in blue and red, respectively. (**D**) Summary table of theoretical versus observed spectra assignments generated under high-resolution MS2 conditions.

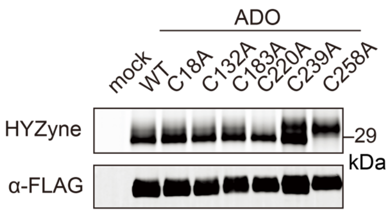

**Fig. S6. HYZyne labelling of Cys mutants of ADO.** HYZyne labeling of wild-type ADO and Cys-to-Ala mutant ADO proteins. Probe labeling (upper) and expression profiles (lower) for HYZyne-treated cells (1 mM, 0.5 h) overexpressing the indicated protein target or Cys mutants thereof.

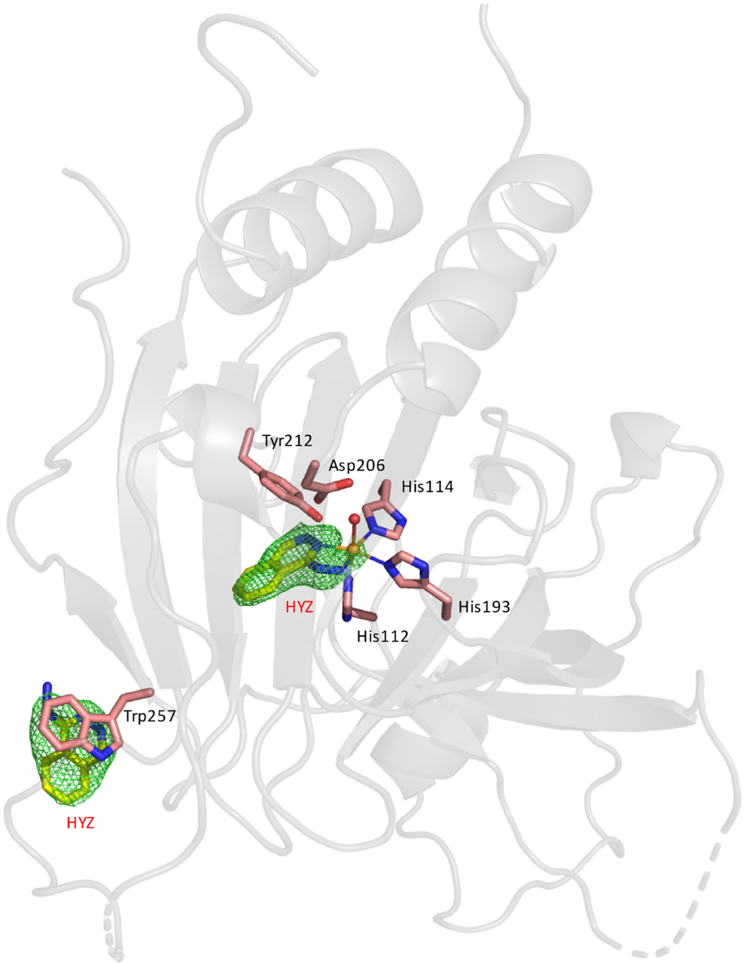

**Fig. S7. Structure of HYZ-bound Co(II)•ADO.** The X-ray crystal structure of HYZ-bound Co(II)•ADO determined at a resolution of 1.88 Å resolution (PDB: 9DMA). The $F_o$–$F_c$ maps are contoured at 3.0 σ and colored in green.

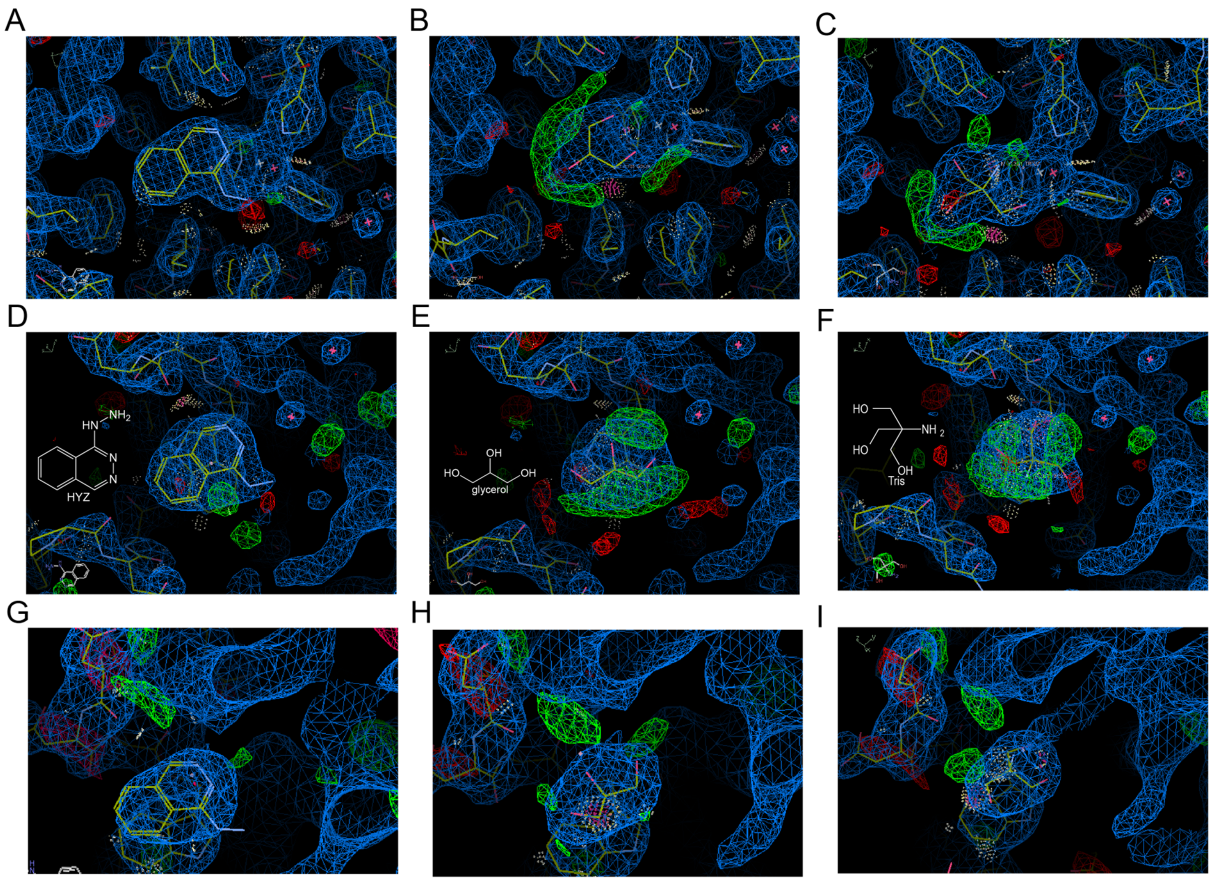

**Fig. S8. Data refinement using small molecules within the crystallization mother liquor.** The 2$F_o$–$F_c$ maps (blue) contoured at 1.5 σ (**A-F**) or 1.2 σ (**G-I**), and the $F_o$–$F_c$ maps (green) contoured at 3.0 σ (**A-I**). (**A-C**) The model and maps using HYZ, glycerol, or Tris in the active sites of Co(II)•ADO (PDB: 9DMA). (**D-F**) The model and maps using HYZ, glycerol, or Tris on the surface of Co(II)•ADO (PDB: 9DMA). (**G-I**) The model and maps using HYZ, glycerol, or Tris on the surface of Fe(II)•ADO (PDB: 9DY4).

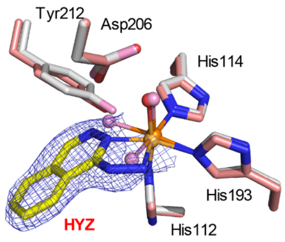

**Fig. S9. Superposition of HYZ-bound Co(II)•ADO (pale orange) and resting-state Co(II)•ADO (gray).** The $2F_o–F_c$ maps are contoured at 1.5 σ and colored in blue. Oxygen atoms of the resting-state Co(II)•ADO are shown in pink, while those of the HYZ-bound Co(II)•ADO are shown in red. HYZ is colored in yellow.

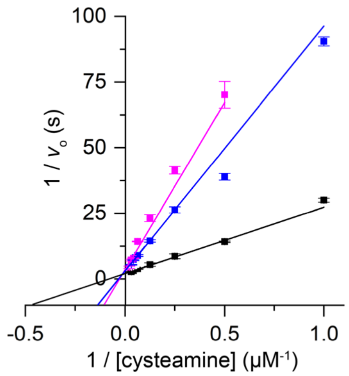

**Fig. S10. Determination of the inhibition pattern of Fe(II)•ADO by HYZ.** Inhibition of 2 μM Fe(II)•ADO was assessed in the presence of varying concentrations of cysteamine and HYZ at 0 (black), 6.67 (blue), and 10 (magenta) μM.

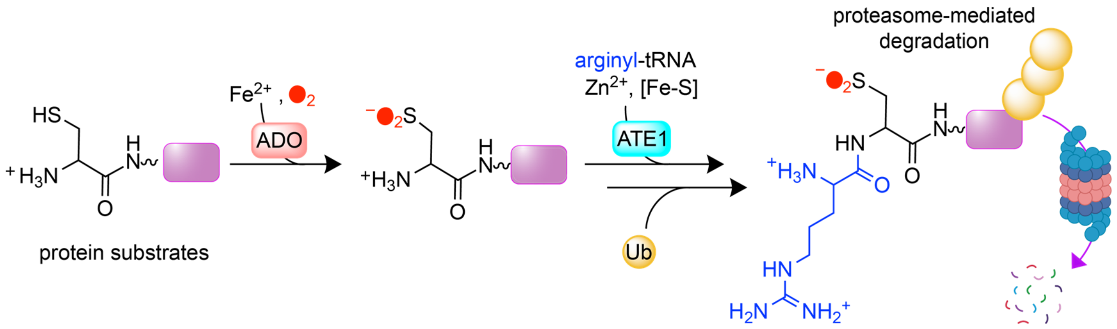

**Fig. S11. *N*-degron pathway scheme.** *N*-degron pathway initiated by *N*-terminal cysteine oxidation via ADO, followed by *N*-terminal arginylation by arginyltransferase (ATE1). Proteins with post-translationally conjugated *N*-terminal Arg residues can undergo ubiquitination and subsequent proteasome-mediated degradation. Ub: ubiquitin. [Part of figure created with BioRender.com]

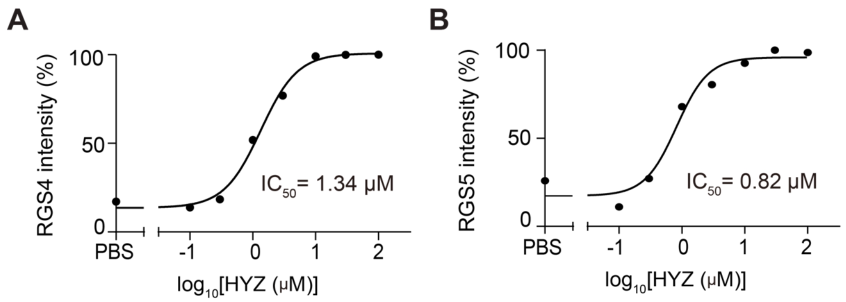

**Fig. S12. Western blot-based inhibition curves of ADO by HYZ in SH-SY5Y cells (1 h treatment).** $IC_{50}$ values were determined by measuring endogenous protein levels of RGS4 (**A**) and RGS5 (**B**) as a function of increasing concentrations of HYZ.

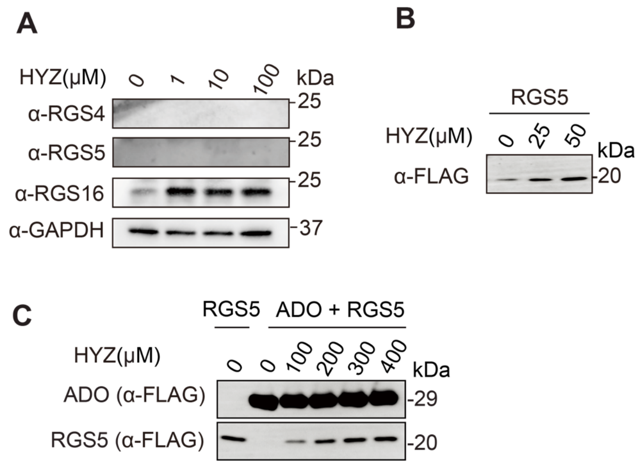

**Fig. S13. ADO inhibition in HEK293T cells.** Inhibition of ADO activity by HYZ in HEK293T cells was quantified with Western blot. (**A**) Endogenous RGS levels in response to different concentrations of HYZ. (**B**) Overexpressed RGS5 bands in response to different concentrations of HYZ. (**C**) Co-overexpressed blots of RGS5 and ADO in the presence or absence of indicated concentrations of HYZ.

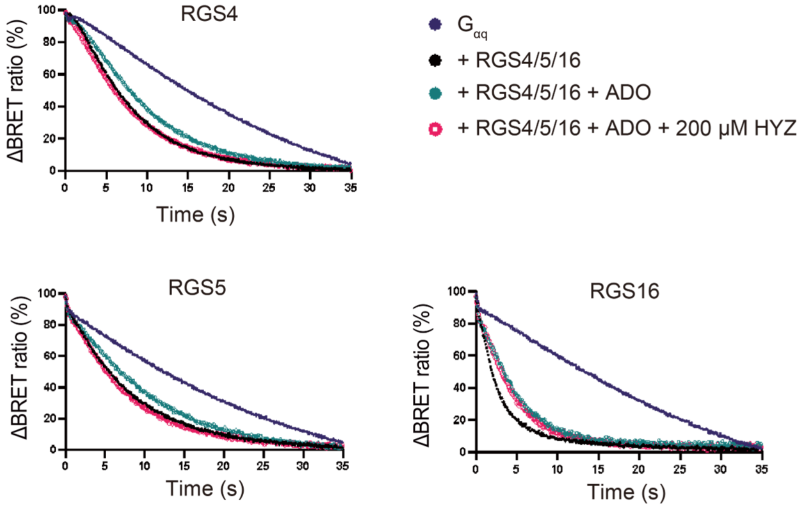

**Fig. S14. BRET assay monitoring the RGS-induced acceleration of G protein deactivation.** Representative traces of BRET signal showing $G_{\alpha q}$ deactivation time course of cells with (black) or without (purple) RGS4/5/16, cells co-transfected with RGS4/5/16 and ADO treated with buffer (green) or 200 μM of HYZ (red). $k_{GAP}$ in Fig. 4D was calculated from ΔBRET.

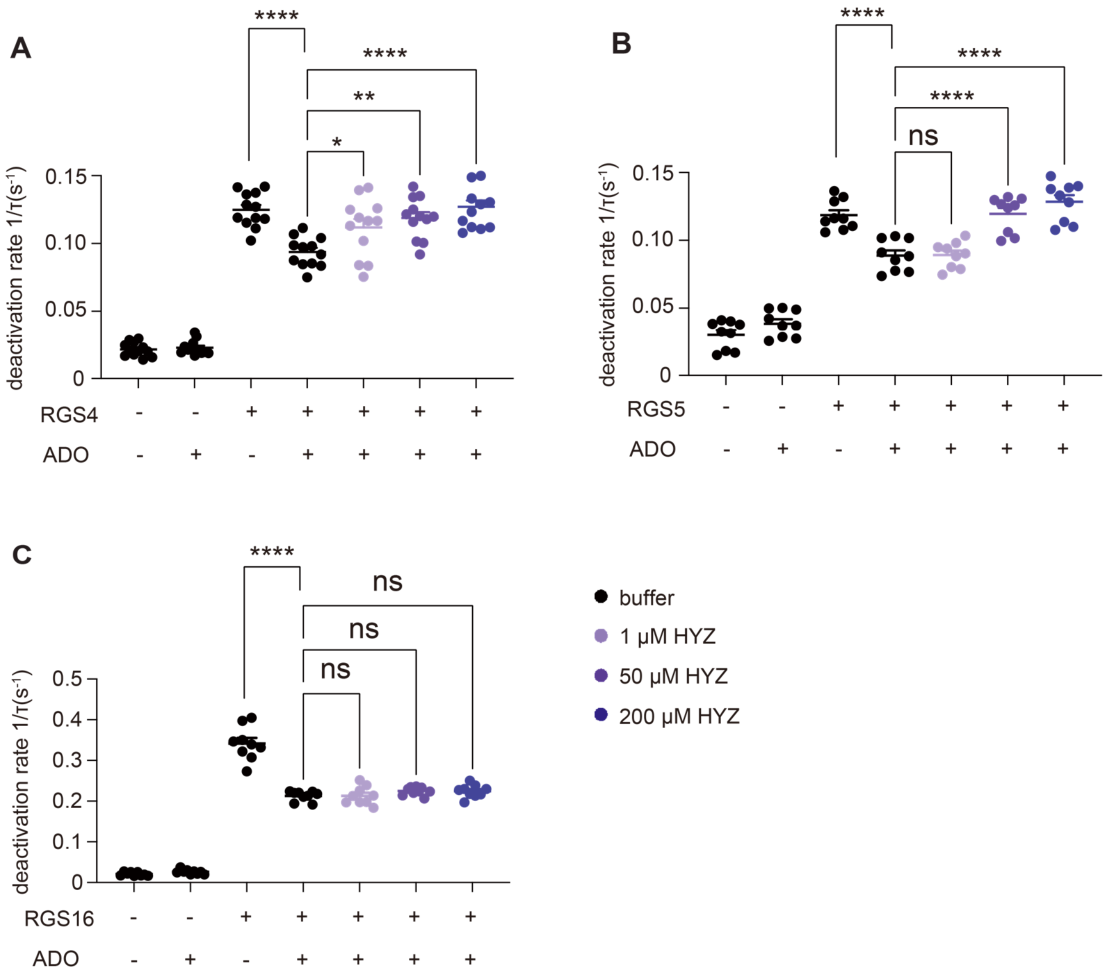

**Fig. S15. HYZ restores the GAP activity of RGS4 and RGS5 in a concentration-dependent manner.** Overexpression of ADO reduces the GAP activity of RGS4 (**A**), RGS5 (**B**), and RGS16 (**C**). Treatment of HYZ concentration-dependently reverses the effect of ADO in RGS4 and RGS5 expressing cells. n = 3 independent experiments with each condition in triplicate. Data represent mean ± SEM. One-way ANOVA with Dunnett's multiple comparisons test (treatments comparison), *$p < 0.05$, **$p < 0.005$, ****$p < 0.0001$, ns = not significant. Unpaired t-test for comparing cells transfected with RGS only and cells co-transfected with RGS and ADO, ***$p < 0.0005$, ****$p < 0.0001$.

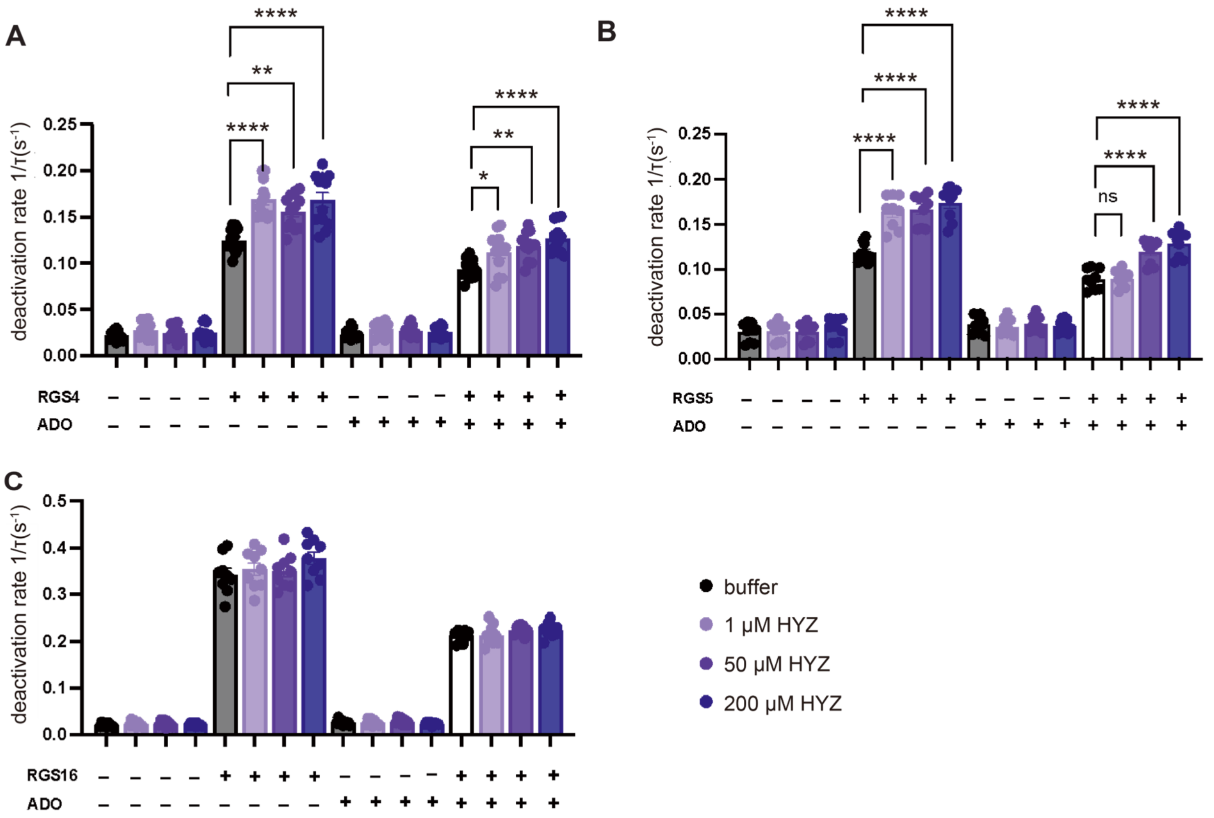

**Fig. S16. Summary of the effect of HYZ on deactivation rate for RGS4/5/16.** GAP activity was measured in the presence or absence of RGS, ADO, and HYZ. Panels show GAP activity for RGS4 (**A**), RGS5 (**B**), and RGS16 (**C**) treated with 0–200 μM HYZ. n = 3 independent experiments with each condition in triplicate. Data represent mean ± SEM. One-way ANOVA with Dunnett's multiple comparisons test, **$p < 0.005$, ****$p < 0.0001$.

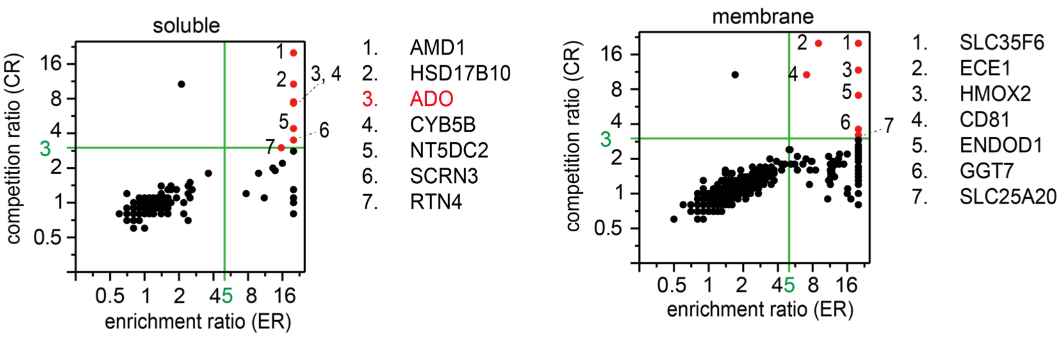

**Fig. S17. Identification of HYZyne targets in U-87 cells.** Quadrant plot of average competition versus average enrichment SILAC protein ratios from quantitative proteomics experiments for soluble and membrane proteomes of U-87 cells. HYZyne (100 μM) and HYZ (100 μM) were used for the ER, while HYZyne (100 μM) and HYZ (500 μM) were used for the CR. Proteins with ER ≥ 5 and CR ≥ 3 (upper right quadrant) were considered high-occupancy targets; listed to the right of the plot. Proteins highlighted red are investigated in this work.

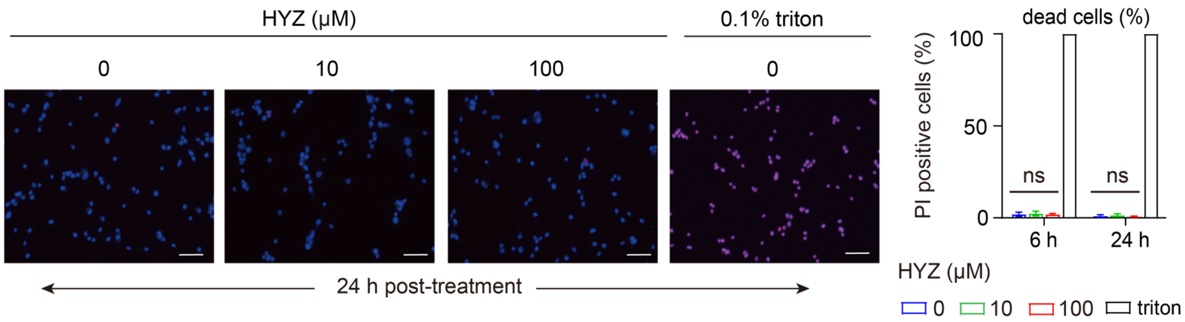

**Fig. S18. Cytotoxicity under HYZ treatment**. Left: Representative images of Hoechst 33342 (blue) and propidium iodide (PI, red) double staining in U-87 cells after 24 h of HYZ treatment. Bar: 200 μm. Right: percentage of PI positive cells at 6 and 24 h of post treatment. n = 3, mean ± SD. Statistical analysis was conducted using one-way ANOVA followed by Tukey's HSD, with each column compared to the untreated HYZ control. The 0.1% triton-treated positive control was excluded from statistical analysis. ns = not significant.

### Table S1. Statistics of monoisotopic masses in MS1 spectra for chemically modified peptides.

| *protein* | *captured peptide* | *tag* | *parent ion intensity* | *charge* | *m/z*$_{theor.}$ | *average m/z*$_{meas.}$ | *error (ppm)* | *SD* | *n* |
|---|---|---|---|---|---|---|---|---|---|
| ADO | SGTSIPL**H**$_{112}$*DHPGMHGMLK | $TEV_{heavy}$ | 1.0E+07 | 3 | 834.4155 | 834.4151 | -0.4 | 0.0003 | 82† |
| | | $TEV_{light}$ | 8.7E+06 | 3 | 832.4109 | 832.4105 | -0.4 | 0.0004 | 84† |
| | QA**C**$_{18}$*LTFR | $TEV_{heavy}$ | 2.6E+08 | 3 | 475.5797 | 475.5793 | -0.7 | 0.0002 | 104† |
| | | $TEV_{light}$ | 2.7E+08 | 3 | 473.5751 | 473.5749 | -0.4 | 0.0003 | 97† |
| | IS**C**$_{132}$*MDK | $TEV_{heavy}$ | 2.7E+07 | 2 | 641.8067 | 641.8065 | -0.3 | 0.0002 | 43† |
| | | $TEV_{light}$ | 2.7E+07 | 2 | 638.7998 | 638.7996 | -0.2 | 0.0003 | 46† |
| | AEYTEASGP**C**$_{183}$*ILTPHR | $TEV_{heavy}$ | 1.3E+07 | 4 | 583.5387 | 583.5383 | -0.5 | 0.0007 | 47† |
| | | $TEV_{light}$ | 1.3E+07 | 4 | 582.0352 | 582.0349 | -0.5 | 0.0005 | 44† |
| | DNLHQIDAVEGPAAFLDILAPPYDPDDGRD**C**$_{220}$*HYYR | $TEV_{heavy}$ | 7.2E+07 | 5 | 912.4318 | 912.4306 | -1.2 | 0.0009 | 97† |
| | | $TEV_{light}$ | 4.1E+07 | 5 | 911.2291 | 911.2281 | -1.0 | 0.0012 | 93† |
| | EASSSA**C**$_{249}$*DLPR | $TEV_{heavy}$ | 2.7E+08 | 3 | 574.6067 | 574.6061 | -0.9 | 0.0004 | 92† |
| | | $TEV_{light}$ | 3.7E+08 | 3 | 572.6021 | 572.6017 | -0.5 | 0.0004 | 91† |
| | EVWLLETPQADDFW**C**$_{258}$*EPYPGPK | $TEV_{heavy}$ | 1.0E+07 | 3 | 1131.5286 | 1131.5271 | -1.3 | 0.0009 | 70† |
| | | $TEV_{light}$ | 9.7E+07 | 3 | 1129.5240 | 1129.5227 | -1.0 | 0.0012 | 69† |

* site of modification by probe clicked with TEV tags.
† the sum of spectral counts (n ≥ 43) for given peptide from two technical replicates.

**Table S2. X-ray crystallographic data collection and refinement statistics**

| | HYZ bound Co(II)•ADO | Fe(II)•ADO soaked in HYZ |
|---|---|---|
| **Data Collection** | SSRL | SSRL |
| Wavelength (Å) | 0.97145 | 0.97946 |
| Space group | $C222_1$ | $C222_1$ |
| Cell dimensions | | |
| a, b, c (Å) | 56.3, 95.1, 117.2 | 56.2, 95.2, 117.8 |
| α, β, γ (°) | 90, 90, 90 | 90, 90, 90 |
| Resolution[a] (Å) | 50.00 – 1.89 | 50.00 – 2.32 |
| | (1.92 – 1.89) | (2.36 – 2.32) |
| Total reflection | 206106 | 104817 |
| Unique reflection | 24951 | 13922 |
| Redundancy | 8.3 (6.4) | 7.5 (5.4) |
| $R_{sym}$ or $R_{merge}$[b] (%) | 10.2 (77.9) | 12.9 (80.6) |
| I/σI | 8.1 (1.02) | 17.4 (1.12) |
| Completeness (%) | 96.8 (92.3) | 99.1 (97.8) |
| $CC_{1/2}$ | 0.999 (0.517) | 0.988 (0.866) |
| **Refinement**[c] | | |
| Resolution (Å) | 48.47 – 1.88 | 48.39 – 2.39 |
| No. reflections | 24745 | 12503 |
| $R_{work}$[d]/$R_{free}$[e] (%) | 21.16 / 25.58 | 25.22 / 30.00 |
| No. Atoms / *B*-factors (Å$^2$) | | |
| Protein | 1905 / 42.0 | 1829 / 85.4 |
| Co/Fe | 1 / 30.9 | 1 / 74.4 |
| HYZ | 24 / 39.1 | 12 / 86.0 |
| Glycerol | 18 / 51.6 | 12 / 88.4 |
| Sulfate | 5 / 72.6 | N/A |
| Water | 100 / 41.9 | 18 / 85.0 |
| r.m.s. deviations | | |
| Bond lengths[d] (Å) | 0.006 | 0.008 |
| Bond angles (°) | 0.867 | 1.090 |
| Ramachandran[e] | | |
| Favored (%) | 98.71 | 96.00 |
| Allowed (%) | 1.29 | 3.60 |
| Outlier (%) | 0.00 | 0.40 |
| **PDB Entry Code** | **9DMA** | **9DY4** |

[a] Values in parentheses are for the highest resolution shell.
[b] $R_{merge} = \Sigma_{hkl}\Sigma_i |I_i(hkl) - \langle I(hkl)\rangle| / \Sigma_{hkl}\Sigma_i I_i(hkl)$, in which the sum is over all the *I* measured reflections with equivalent miller indices *hkl*; $\langle I(hkl)\rangle$ is the averaged intensity of these *i* reflections, and the grand sum is over all measured reflections in the data set.
[c] All positive reflections were used in the refinement.
[d] According to previous literature(*59*).
[e] Ramachandran statistics were analyzed using MolProbity(*60*). The outlier in 9DY4 is Pro115.

**Table S3. Comparison of the EPR parameters for Co(II)•ADO, Co(II)•ADO (cysteamine), and Co(II)•ADO (HYZ)**

| Sample | *S* | *g* values ($g_x$, $g_y$, $g_z$) |
|---|---|---|
| Co(II)•ADO(*24*) | 3/2 | 2.63, 4.25, 5.26 |
| Co(II)•ADO (cysteamine)(*24*) | 3/2 | 2.25, 4.00, 5.71 |
| Co(II)•ADO (HYZ) | 3/2 | 2.55, 4.41, 5.40 |

**Data S1.** Compiled average SILAC protein ratios with standard deviation for enrichment and competition experiments with HYZyne for HEK293T and U-87 cell lines followed by representative datasets for each type of experiment. These datasets are shown in separate tabs (8 in total), each displaying median SILAC ratios for all quantified tryptic peptides per protein. Corresponding peptide sequences, masses, charge states, and individual SILAC ratios are shown. Additional datasets are available upon request.

**Data S2.** Compiled average SILAC and ReDiMe protein ratios with standard deviation for enrichment and competition experiments with HYZyne for RAW264.7 and Neuro 2a cell lines, and enrichment experiments with mouse heart and brain followed by representative datasets for each type of experiment. These datasets are shown in separate tabs (12 in total), each displaying median SILAC or ReDiMe ratios for all quantified tryptic peptides per protein. Corresponding peptide sequences, masses, charge states, and individual SILAC or ReDiMe ratios are shown. Additional datasets are available upon request.

**Data S3.** Primers used for qPCR.
